# Proteomic Analysis of Huntington’s Disease Medium Spiny Neurons Identifies Alterations in Lipid Droplets

**DOI:** 10.1101/2022.05.11.491152

**Authors:** Kizito-Tshitoko Tshilenge, Carlos Galicia Aguirre, Joanna Bons, Nathan Basisty, Sicheng Song, Jacob Rose, Alejandro Lopez-Ramirez, Akos Gerencser, Swati Naphade, Ashley Loureiro, Cameron Wehrfritz, Anja Holtz, Sean Mooney, Birgit Schilling, Lisa M. Ellerby

**Affiliations:** The Buck Institute for Research on Aging, Novato, California, 94945, USA; University of Southern California, Leonard Davis School of Gerontology, 3715 McClintock Ave, Los Angeles, CA 90893, USA; Translational Gerontology Branch, National Institute on Aging (NIA), NIH, Baltimore, Maryland, 21244, USA; Department of Biomedical Informatics and Medical Education, School of Medicine, University of Washington, Seattle, WA, 98109, USA

**Author notes:** Correspondence: Birgit Schilling and Lisa M. Ellerby. Abbreviations: AD, Alzheimer’s disease; ApoE, apolipoprotein E; BDNF, brain-derived neurotrophic factor; CAG, cytosine adenine guanine; CNS, central nervous system; CREB, cAMP response element-binding protein; CV, compensation voltage; DDA, data-dependent acquisition; DE, differentially express; DIA, data-independent acquisition; ECM, extracellular matrix; EMT, epithelial-mesenchymal transition; FAIMS, high-field asymmetric waveform ion mobility spectrometry; FDR, false discovery rate; GABA, gamma aminobutyric acid; GO, Gene Ontology; GPX, glutathione peroxidase; HD, Huntington’s disease; HTT, Huntingtin; IGFBP7, insulin-like growth factor-binding protein 7; IFN-γ, interferon-gamma; IPA, Ingenuity Pathway Analysis; iPSC, induced pluripotent stem cell; LC-MS/MS, liquid chromatography-tandem mass spectrometry; LFQ, label-free quantitation; LGE, lateral ganglionic eminence; logFC, log fold change; MCM, minichromosome maintenance; MHC, major histocompatibility complex; mHTT, mutant Huntingtin; MS, mass spectrometry; MSN, medium spiny neurons; NSC, neural stem cells; PBS, phosphate buffered saline; PD, Parkinson’s disease; polyQ, polyglutamine; SASP, senescence-associated secretory phenotype; SMA, spinal muscular atrophy; STMN1, stathmin-1; TrkB, tyrosine kinase receptor B; WNT, wingless Int-1; XIC, extracted ion chromatogram.

**Keywords:** Neurodegeneration, Huntington’s disease, induced pluripotent stem cells, medium spiny neurons, quantitative proteomics, data-independent acquisitions, data-dependent acquisitions, ion mobility.

## Abstract

Huntington’s disease (HD) is a neurodegenerative disease caused by a CAG repeat expansion in the Huntingtin (*HTT*) gene. The resulting polyglutamine (polyQ) tract alters the function of the HTT protein. Although HTT is expressed in different tissues, the medium spiny projection neurons (MSNs) in the striatum are particularly vulnerable in HD. Thus, we sought to define the proteome of human HD patient-derived MSNs. We differentiated HD72 induced pluripotent stem cells and isogenic controls into MSNs and carried out quantitative proteomic analysis by two approaches. First, using data-dependent acquisitions with FAIMS (FAIMS-DDA) for label-free quantification on the Orbitrap Lumos mass spectrometer, we identified 6,323 proteins with at least two unique peptides (FDR ≤ 0.01). Of these, 901 proteins were significantly altered in the HD72-MSNs, compared to isogenic controls. Second, we quantitatively validated protein candidates by comprehensive data-independent acquisitions on a TripleTOF 6600 mass spectrometer quantifying 3,106 proteins with at least two unique peptides. Functional enrichment analysis identified pathways related to the extracellular matrix, including TGF-β regulation of extracellular matrix, epithelial-mesenchymal transition, DNA replication, senescence, cardiovascular system, organism development, regulation of cell migration and locomotion, aminoglycan glycosaminoglycan proteoglycan, growth factor stimulus and fatty acid processes. Conversely, processes associated with the downregulated proteins included neurogenesis-axogenesis, the brain-derived neurotrophic factor-signaling pathway, Ephrin-A: EphA pathway, regulation of synaptic plasticity, triglyceride homeostasis cholesterol, plasmid lipoprotein particle immune response, interferon-γ signaling, immune system major histocompatibility complex, lipid metabolism and cellular response to stimulus. Moreover, proteins involved in the formation and maintenance of axons, dendrites, and synapses (e.g., Septin protein members) are dysregulated in HD72-MSNs. Importantly, lipid metabolism pathways were altered, and we found that lipid droplets accumulated in the HD72-MSNs, suggesting a deficit in lipophagy. Our proteomics analysis of HD72-MSNs identified relevant pathways that are altered in MSNs and confirm current and new therapeutic targets for HD.

## INTRODUCTION

Huntington’s disease (HD) is a rare progressive monogenic neurological disorder caused by a trinucleotide repeat expansion in exon-1 of the Huntingtin gene (*HTT*) (1–3). The clinical hallmark of HD is a chorea that co-exists with cognitive decline and emotional disturbances (4). There is no cure, and no treatment alters the course of this devasting disease. HD phenotypes are linked to expression of mutant HTT protein (mHTT) that harbors expanded glutamine stretches (over 38) in the N-terminal region. HD features neuronal degeneration in the brain, and medium-spiny projections neurons (MSNs) within the striatum are particularly vulnerable (5).

While substantial progress has been made towards elucidating how the CAG repeats within *mHTT* lead to the clinical outcomes in HD, our understanding of the mechanisms underlying the motor deficits and striatal degeneration is incomplete (6). Those mechanisms likely occur in parallel. For example, mHTT has altered localization, conformation, and protein interactions (7–11). Proteolysis of mHTT generates N-terminal fragments containing the polyQ expansion is found in HD human brain and mouse models (9, 12–18). The cleaved forms of the protein are found in multiple cellular compartments, including the nucleus, and cause aberrant interactions of mHTT with key partners, such as transcription factors, autophagy and mitochondrial proteins, and thus lead to neuronal death in the striatum and cortex (19–21). Deciphering how the mHTT alters the proteome is critical to understanding HD molecular mechanisms, and few studies have focused on defining the proteomes of human HD patient-derived neurons.

Recent progress in mass spectrometry (MS)-based proteomics has allowed significant improvements in proteome resolution, sensitivity and depth-coverage of a given biological system (22, 23). Previous studies used quantitative MS-based proteomics to measure relative changes in the protein abundances in human postmortem HD frontal cortex and identified signaling pathways that are dysregulated in HD, including Rho-mediated, actin cytoskeleton and integrin signaling, mitochondrial dysfunction and axonal guidance (24). Comprehensive quantitative proteomics applied to investigate spatiotemporal mechanisms of mHtt in R6/2 HD mice characterized the insoluble proteome during disease progression and highlighted extensive dysregulation in brain regions vulnerable to HD (25). A recent review highlights the proteomics carried out in HD and some of the critical gaps in the field, including the generation of robust human HD cell-type-specific proteome data sets (26).

Disease modeling in human induced pluripotent stem cells (iPSCs) allows central nervous system (CNS)-relevant cells to be generated *in vitro* and molecular defects to be identified that contribute to polyQ-expansion disorders (27–30). In our previous work, we used human patient– derived HD-iPSCs (72CAG/19CAG, HD72) and genetically corrected the cells to a normal repeat length (21CAG/19CAG, C116), thus creating an isogenic control (31). Our group showed that HD phenotypes manifest in differentiated neural stem cells (NSCs), not in iPSCs (32, 33). Further, our transcriptomic analysis of isogenic HD72-NSCs suggested that HD is linked to developmental impairments that prevent the proper generation of MSNs and subsequent loss of MSN identity (27, 31, 32, 34, 35). So far, quantitative proteomics of iPSCs modeling HD focused on undifferentiated stem cells or unrestricted neuronal populations (36, 37). In contrast, the proteome in directed HD72-MSNs derived from iPSCs has not been explored.

To define the proteomic signature in isogenic HD72-MSNs and to determine how mHTT leads to neurotoxicity in MSNs, we performed comprehensive quantitative proteomics by liquid chromatography-tandem mass spectrometry (LC-MS/MS) using complementary approaches. First, we compared triplicates of HD72-MSNs and isogenic controls in which the CAG expansion was genetically corrected to a normal repeat length. We used an unbiased discovery workflow combining a modern high-field asymmetric waveform ion mobility spectrometry (FAIMS) device (38) with an Orbitrap Lumos mass spectrometer operating in data-dependent acquisition (DDA) mode for the identification and label-free quantification MS1-based (LFQ) of significantly changing protein candidates (Discovery). Subsequently, we acquired the same samples by data-independent acquisition (DIA) on a TripleTOF 6600 mass spectrometer for further quantification and validation (MS2-based Quantification, DIA) (39). Our proteomic workflow is summarized in **Fig. 1**.

**Figure 1.**
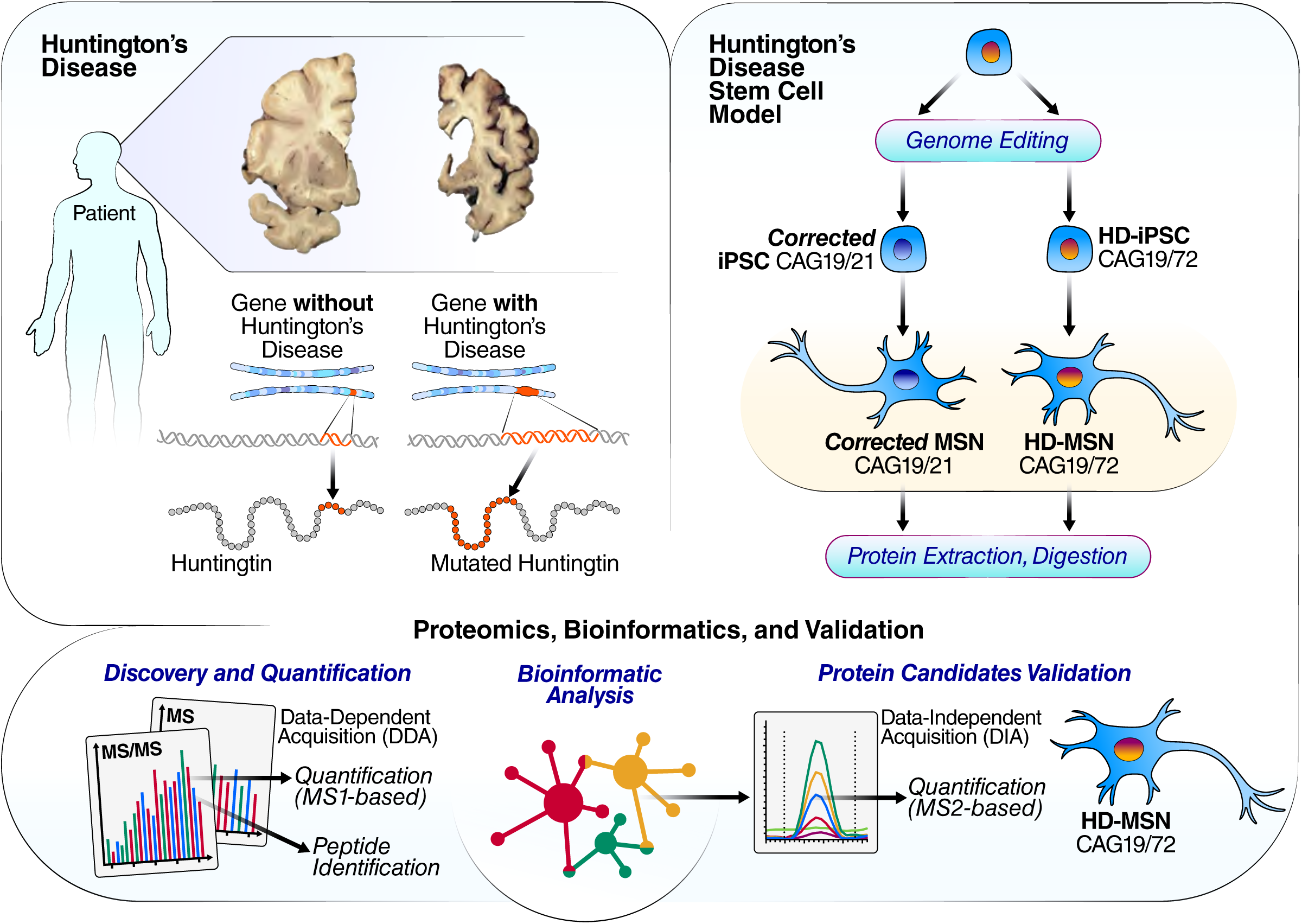
Schematic representation of HD, isogenic HD-MSN and proteomics workflow. HD is a monogenic disease caused by a CAG (coding for glutamine) expansion in the HTT gene. The striatum, depicted in the upper left panel, is heavily affected in HD. The inhibitory medium spiny neurons are lost during disease progression. Total proteins were isolated from C116-MSN and HD72-MSN cultures (in triplicates). Samples were digested and subjected to a comprehensive quantitative proteomic analysis with deep coverage with FAIMS ion mobility separation coupled to data-dependent acquisition mode on an Orbitrap Lumos mass spectrometer for label-free quantification. Subsequently, significantly changed proteins were validated with an independent quantitative approach collecting data-independent acquisitions on a TripleTOF 6600 mass spectrometer. After protein identification and quantification, bioinformatic analyses were used to identify molecular pathways and networks relevant in HD.

The FAIMS-DDA MS workflow enabled high-resolution and high-sensitivity acquisitions, which, when combined with the additional ion mobility dimension, allowed us to achieve deep proteome coverage (6,323 proteins). Functional enrichment analysis of the upregulated proteins identified pathways related to the extracellular matrix (ECM), epithelial-mesenchymal transition (EMT), DNA replication, senescence, cardiovascular system, organism development, regulation of cell migration and locomotion, aminoglycan glycosaminoglycan proteoglycan, growth factor stimulus and fatty acid processes. Conversely, processes associated with the downregulated proteins include neurogenesis-axogenesis and, more specifically, the brain-derived neurotrophic factor-signaling pathway, Ephrin-A:EphA pathway, regulation of synaptic plasticity, triglyceride homeostasis cholesterol, plasmid lipoprotein particle immune response, interferon-γ signaling, immune system major histocompatibility complex (MHC), lipid metabolism and cellular response to stimulus. We validated the role of DNA and lipid signaling alterations in HD-MSNs and showed that lipid droplets accumulate in an HD phenotype. Overall, our proteomics analysis of isogenic HD72-MSNs provides a comprehensive catalog of proteins that play a role in MSN cellular phenotypes and confirm current and new therapeutic targets for HD.

## EXPERIMENTAL PROCEDURES

### Human Induced Pluripotent Stem Cell-Derived NSC Culture

C116 and HD72 iPSCs were maintained in mTeSR™1 (STEMCELL Technology, 05850) medium before the differentiation. To induce iPSCs toward a neuroepithelial fate, we used a monolayer differentiation approach with modifications. Briefly, iPSCs were manually cleaned by removing any colonies with spontaneous differentiation. To initiate differentiation (Day 0), iPSCs were passaged with 1 mg/mL collagenase [(Type IV, Thermo Fisher Scientific, 17104019) in Gibco KnockOut DMEM/F-12 medium (Thermo Fisher Scientific, 12660012)] for 35 min at 37°C. The colonies were gently detached by scraping, and the cell aggregates were triturated by pipetting 2–3 times with a 2-mL pipette to yield a uniform suspension of aggregates and avoiding creating a single-cell suspension. The cell aggregates were transferred onto a Matrigel (1 mL, 50 µg, Corning, CB-40234)-coated plate containing mTeSR™1 and incubated at 37°C. For neural induction (Day 2), SMAD signaling was inhibited by adding SB431542 (10 µM, Tocris, 1614) and LDN-193189 (1 µM, Tocris, 6053) in mTeSR™1 during the medium changing, thus promoting neuroectodermal differentiation and suppressing mesoderm and endoderm fates (70). From day 4, the mTeSR™1 medium was changed every day in presence of SB431542 (10 µM, Tocris, 1614) and LDN-193189 (1 µM, Tocris, 6053). From day 8, the colonies became organized and increased in size. The center of colonies became dense and compact, and the peripheral regions presented elongated cells. At day 10, the colonies were first cleaned to remove peripheral regions, and then the dense center regions were manually picked by scraping after a collagenase treatment of 25 min at 37°C, to avoid over-collagenase. Cell aggregates from the center regions were transferred at low density to minimize merging, into a 10-cm Matrigel (1 mL, 50 µg, Corning, CB-40234)-coated plate containing N2B27 medium [(DMEM/F12, Gibco, Thermo Fisher Scientific, 11320-033) supplemented with 1 X N2 (Thermo Fisher Scientific, 17502001), 1 X B27 (Thermo Fisher Scientific, 17504001), 1 X GlutaMAX (Thermo Fisher Scientific, 35050061), 1 X Non-Essential Amino Acids (Thermo Fisher Scientific, 11140050), 25 ng/mL β-FGF (Peprotech, 100-18B) and 100 U/mL penicillin-streptomycin (Thermo Fisher Scientific, 15140122)]. The cells were cultured in presence of 25 ng/mL Activin A (PeproTech, AF-120-14E) to induce regional patterning toward a lateral ganglionic eminence (LGE) identity (37). The N2B27 medium was changed every 2 d in presence of Activin A and β-FGF. At day 12, neuroepithelial differentiation became apparent with the formation of small neural rosettes showing a columnar shape that further organized and increased in size at day 14 in forming neural tube-like structures with a central lumen and three-dimensional growth. Thereafter, the neural rosette structures were mechanically selected by separating the island from the surrounding cells with a needle to minimize contamination with non-neural cells. The isolated rosettes were triturated by pipetting 8–10 times with 1,000-µl pipette tip. At least 15–20 neural rosettes/well were plated in a Matrigel-coated P12 well plate in presence of Neural Proliferation Medium [Neurobasal medium (Thermo Fisher Scientific, 21103049), B27-supplement 1 X (Thermo Fisher Scientific, 17504001), GlutaMAX 1 X (Thermo Fisher Scientific, 35050061), 10 ng/mL leukemia inhibitory factor (PeproTech, 300-05), 100 U/mL penicillin-streptomycin] supplemented with 25 ng/mL β-FGF and 25 ng/mL Activin A. The resulting neural stem cells (NSCs) were passaged when cell cultures became confluent. The passaging cells were moved gradually from P6, P12 wells to 6-cm plates with the cells plated at a high density. Nestin, SOX1, SOX2, and PAX6 staining of NSCs validated the cell type.

### MSN Differentiation

Activin A (25 ng/mL, PeproTech, AF-120-14E)-generated C116 and HD72 NSCs were used to prepare MSNs. Nunc six-well plates were treated with poly-D-lysine hydrobromide (1 mL, 100 µg/mL by Sigma Aldrich, P6407) and incubated (37°C and 5% CO2) overnight (ON). Corning cell culture grade water, 25-055-CVC (1 mL), was used to wash plates and the plates were dried for 1 h. Next, the plates were treated with Matrigel (1 mL, 50 µg, Corning, CB-40234) overnight in a 37°C incubator. MSNs were prepared according to Kemp et al. (30). Synaptojuice A medium (2 mL) was used for seeding NSCs (1x10^6^ per well). Synaptojuice A was prepared with 10X synaptojuice A supplement (5 mL), advanced DMEM/F12 medium (44.1 mL, Gibco, 12634010), penicillin/streptomycin (P/S) (450 µL, Invitrogen, 15140122), and 100X Glutamax (450 µL, Invitrogen, 35050079). Synaptojuice A supplement (10X) contains advanced DMEM/F12 medium (38 mL, Thermo Fisher Scientific, 12634010), MACS NeuroBrew-21 with retinoic acid with final concentrations noted (10 mL, MACS Miltenyi Biotec, 130-093-566), PD0332991 (20 µM, Tocris Bioscience, 4786), DAPT (100 µM, Tocris Bioscience, 2634), human BDNF (100 ng/mL, MACS Miltenyi Biotec, 130-096-286), LM22A4 (5 µM, Tocris Biotec, 4607), forskolin (100 µM, Tocris Bioscience, 1099), CHIR 99021 (30 µM, Tocris Bioscience, 1099), GABA (3 mM, Tocris Bioscience, 0344), CaCl2 (1.8 mM, Tocris Bioscience, 3148), ascorbic acid (2 mM, Tocris Bioscience, 4055). Medium was passed through a 0.22-µm filter. Cells were treated with synaptojuice A (2 mL) for 7 days, performing half-medium changes every other day. On day 8, full medium changes were completed and then, the cells were treated with synaptojuice B (2 mL) for the next 7 days. Synaptojuice B was prepared with 10X synaptojuice B (5 mL) supplement and basal medium (45 mL). Basal medium contains advanced DMEM/F12 medium (22.5 mL,

Thermo Fisher Scientific, 12634010), penicillin/streptomycin (P/S) (450 µL, Invitrogen, 15140122), and 100X Glutamax (450 µL, Invitrogen, 35050079) and Neurobasal A Medium (22.5 mL, Gibco, 10888022), penicillin/streptomycin (P/S) (450 µL, Invitrogen, 15140122), and 100X Glutamax (450 µL, Invitrogen, 35050079). Synaptojuice B supplement contains advanced DMEM/F12 medium (19.7 mL, Thermo Fisher Scientific, 12634010), Neurobasal A medium (19.7 mL, Gibco, 10888022), MACS NeuroBrew-21 with retinoic (10 mL, MACS Miltenyi Biotec, 130- 093-566), PD0332991 (100 µL, 20 µM, Tocris Bioscience, 4786), human BDNF (50 µL, 100 ng/mL, MACS Miltenyi Biotec, 130-096-286), LM22A4 (25 µL, 5 µM, Tocris Biotec, 4607), CHIR 99021 (250 µL, 30 µM, Tocris Bioscience, 1099), GABA (500 µL, 3 mM, Tocris Bioscience, 0344), CaCl2 (370 µL, 1.8 mM, Tocris Bioscience, 3148) and ascorbic acid (100 µL, 2 mM, Tocris Bioscience, 4055). Synaptojuice B medium was filtered through a 0.22-µm filter. Cells were treated with synaptojuice B (2 mL) for 7 days, and half medium changes were performed until day 14.

### MSNs Treatments with IFN-γ

For the IFN-γ experiments, prepatterned Activin A–treated NSCs from C116 and HD72 were plated at 90,000 cells per well in an eight-well chamber slide for MSN differentiation. After synaptojuice A and B treatment, MSNs were stimulated for 48 h with IFN-γ (PeproTech, 300-02- 100UG) at different concentrations: 10, 50, 100 and 200 ng/mL. Non-treated MSNs were used as control. For each treatment, a duplicate was performed.

### Cell Immunofluorescence of Human MSNs

Cells were fixed using 4% paraformaldehyde (Sigma, 158127) in 0.1 M phosphate-buffer saline (PBS), pH 7.4 (Corning, 21-040-CV) for 30 min. After three washes in cold PBS, cells were permeabilized and blocked for 1 h at RT using 0.1% Triton X-100 (Thermo Fisher Scientific, 28313) and 4% normal donkey serum (Jackson Immuno Research, 017-000-121) in PBS. Primary antibodies were added in the presence of blocking buffer overnight at 4°C. Secondary antibodies (1:500) were added after three PBS washes in blocking buffer at RT for 1 h. The following primary antibodies were used for the immunofluorescence studies: rabbit anti-DARPP-32 (Santa Cruz, sc-271111, 1:100), rabbit anti-MAP2 (Millipore, AB5622, 1:100), rabbit anti-Nestin (Abcam, ab92391, 1:00), mouse anti-MHC-class-II (Abcam, ab55152, 1:100), rabbit anti-Cleaved Caspase-3 (CellSignal, 9661, 1:100) and mouse anti-HTT (Millipore, MAB2166, 1:100). The secondary antibodies were donkey anti-rabbit, anti-mouse IgG conjugated with Alexa-546 (Invitrogen, A10040 and A10036) or Alexa-647 (Invitrogen, A-31573 and A-31571). Images were acquired using a Biotek Cytation 5 microscope and were prepared using Fiji software (ImageJ).

### Protein Extraction for Proteomic Analysis

Triplicate samples of cultured C116-MSNs and HD72-MSNs were washed three times with cold PBS 1X, pH 7.4 (Corning, 21-040-CV), and total proteins were isolated using 300 µl of cold mammalian protein extracting reagent (Thermo Fisher Scientific, 78501) containing protease inhibitor cocktail (cOmplete, Mini Protease Inhibitor Cocktail, Roche, 11836170001). The cell lysate was harvested by scraping and transferred directly into a cold 1.5-mL tube and stored at - 80°C.

### Proteomic Sample Preparation

#### Chemicals

Acetonitrile (AH015) and water (AH365) were from Burdick & Jackson (Muskegon, MI). Iodoacetamide (I1149), dithiothreitol (DTT, D9779), formic acid (94318-50ML-F), and triethylammonium bicarbonate buffer 1.0 M, pH 8.5 (T7408) were from Sigma Aldrich (St. Louis, MO), urea (29700) was from Thermo Scientific (Waltham, MA), sequencing grade trypsin (V5113) was from Promega (San Luis Obispo, CA), and HLB Oasis SPE cartridges (186003908) were from Waters (Milford, MA).

#### Protein Precipitation, Digestion and Desalting

Protein samples were precipitated with a ProteoExtract Protein Precipitation Kit (539180) from MilliporeSigma (Burlington, MA) as per the manufacturer’s protocol. Samples were resuspended in 50 mM triethylammonium bicarbonate. Total protein concentration was determined with a BCA kit (23227) from Thermo Fisher (Waltham, MA). Aliquots of each sample containing ∼100 μg of protein were brought to equal volumes with 50 mM triethylammonium bicarbonate buffer at pH 8. The mixtures were reduced with 20 mM DTT (37 °C for 1 h) and then alkylated with 40 mM iodoacetamide (30 min at RT in the dark). Samples were diluted 10-fold with 50 mM triethylammonium bicarbonate buffer at pH 8 and incubated overnight at 37°C with sequencing grade trypsin (Promega, San Luis Obispo, CA) at a 1:50 enzyme:substrate ratio (wt/wt). Peptide supernatants were collected and desalted with Oasis HLB 30-mg Sorbent Cartridges (Waters, Milford, MA; 186003908), concentrated, and resuspended in a solution containing mass spectrometric “Hyper Reaction Monitoring” retention time peptide standards (HRM, Biognosys, Schlieren, Switzerland; Kit-3003) and 0.2% formic acid in water.

### Mass Spectrometric Analysis

#### Orbitrap Lumos FAIMS DDA MS Analysis

Triplicate samples from corrected C116-MSNs and HD-MSNs were analyzed by reverse-phase HPLC-ESI-MS/MS on the EASY-nLC 1200 system and analytical column (Thermo EASYspray 50 cm x 75 µm ID, PepMap C18 2 µm, 100 Å) coupled to the Orbitrap Lumos mass spectrometer (Thermo Fisher Scientific, San Jose, CA) with an EASY-Spray source. Column temperature was set to 50°C. Mobile phase A was 0.1% formic acid in water, and mobile phase B was 0.1% formic acid in 80% acetonitrile and 19.9% water. Flow rate was set at 300 nL/min, and a two-stage gradient was used for each sample: 1) 7–30% mobile phase B over 125 min; and 2) 30–45% mobile phase B over 40 min. For each sample, 2 µg of peptides were injected onto the column. All samples were analyzed by DDA. For DDA analysis, full MS scans were performed over m/z 380–1,580 with the Orbitrap analyzer operating at 240,000 resolution and AGC = 400,000 ions with ‘high-field asymmetric waveform ion mobility spectrometry‘ (FAIMS) settings enabled at three compensation voltages (CVs): -50 V, -65 V, -85 V. Each of the selected CVs was applied to sequential survey scans and MS/MS cycles (1 s per CV). Survey scans were followed by MS2 scans of the most intense precursor ions for 1 s. MS2 scans were performed by 0.7 m/z isolation with the quadrupole, normalized higher-energy collisional dissociation (HCD) collision energy of 35%. Dynamic exclusion was set to 30 s, mass tolerance to 10 ppm, and intensity threshold to 5,000. Maximum injection time was set to 30 ms, AGC target was set to 10,000 ions, charge states +1 or >+8 were excluded, and the advanced peak determination was toggled on.

#### TripleTOF 6600 DIA MS Analysis

Samples were analyzed by reverse-phase HPLC-ESI-MS/MS using the Eksigent Ultra Plus nano-LC 2D HPLC system (Dublin, CA) combined with a cHiPLC system directly connected to an orthogonal quadrupole time-of-flight TripleTOF 6600 mass spectrometer (SCIEX, Redwood City, CA). Typically, mass resolution in precursor scans was approximately 45,000, and fragment ion resolution was approximately 15,000 in “high sensitivity” product ion scan mode. After injection, peptide mixtures were transferred onto a C18 pre-column chip (200 μm × 6 mm ChromXP C18-CL chip, 3 μm, 300 Å; SCIEX) and washed at 2 μL/min for 10 min with the loading solvent (H2O/0.1% formic acid) for desalting. Peptides were transferred to the 75 μm × 15 cm ChromXP C18-CL chip, 3 μm, 300 Å (SCIEX) and eluted at 300 nL/min with a 3-h gradient using aqueous and acetonitrile solvent buffers. All samples were analyzed by DIA, specifically using variable window DIA acquisitions (40). In these DIA acquisitions, 64 windows of variable width (5–90 m/z) were passed in incremental steps over the full mass range (m/z 400– 1,250) with an overlap of 1 m/z. The cycle time of 3.2 seconds included a 250-ms precursor ion scan, followed by acquisition of 64 DIA MS/MS segments, each with a 45-ms accumulation time (see **supplemental Table S1** for the window isolation scheme). The variable windows were determined according to the complexity of the typical MS1 ion current observed within a certain m/z range using a SCIEX “variable window calculator” algorithm (more narrow windows were chosen in “busy” m/z ranges, wide windows in m/z ranges with few eluting precursor ions) (41). DIA tandem mass spectra produce complex MS/MS spectra, which are a composite of all the analytes within each selected Q1 m/z window.

### Data Processing

For FAIMS DDA experiments, data analysis was performed with Proteome Discoverer version 2.3.0.523 (Thermo Fisher Scientific). The database search was performed using SEQUEST HT (Thermo Fisher Scientific), and parameters were as follows: SwissProt human protein database (20,417 entries, 09 April 2019), trypsin enzyme digestion allowing two missed cleavages, 10-ppm precursor ion mass tolerance, and 0.6-Da fragment ion mass tolerance. Dynamic modifications were methionine oxidation (+15.995 Da) and N-terminal protein acetylation (+42.011 Da), and a static modification was defined as cysteine carbamidomethylation (+57.021 Da). Identifications were filtered to 1% false discovery rate (FDR) (PSM, peptide and protein levels) with Percolator (42). Label-free quantification (LFQ) was performed within Proteome Discoverer using razor and unique peptides, and chromatographic alignment was enabled (maximum 10-min retention time shift and 10-ppm mass tolerance). Abundance was normalized to the total peptide amount, and scaled on control average. Modified peptides were excluded from quantification, and peptide quantities were summed for protein abundances. Statistical analysis was performed using ProStaR software suite (43). Proteins with less than two unique peptides and proteins with more than three missing values across all conditions were removed. Data were log2-transformed and missing values were replaced by the 2.5 percentile value for the POV (Partially Observed Values) and MEC (Missing on the Entire Condition). Pairwise protein statistics were performed using a Limma t-test, and an absolute log2(fold-change) threshold set at 0.58. Slim (sliding linear model) method (44) was applied to adjust p-values for multiple testing, and significantly altered proteins were sorted out using a p-value threshold that guarantees a FDR at 1.04%.

For DIA quantification, all collected data were processed in Spectronaut (version 14.2.200619.47784) using Biognosys (BGS) factory settings. Briefly, calibration was set to non-linear iRT calibration with precision iRT selected. DIA data were matched against a panhuman library that provides quantitative DIA assays for 10,316 human proteins (45) and supplemented with scrambled decoys (library size fraction of 0.1), using dynamic mass tolerances and dynamic extraction windows. The DIA/SWATH data was processed for relative quantification comparing peptide peak areas from various different time points during the cell cycle. For the DIA/SWATH MS2 data sets, quantification was based on XICs of 3-6 MS/MS fragment ions, typically y- and b-ions, matching to specific peptides present in the spectral library. Interference correction was enabled on MS1 and MS2 levels. Precursor and protein identifications were filtered to 1% FDR, estimated using the mProphet algorithm (46). Quantification was normalized to local total ion chromatogram. Statistical comparison of relative protein changes was performed with paired t-tests, and p-values were corrected for multiple testing, specifically applying group wise testing corrections using the Storey method (47). Finally, proteins identified with less than two unique peptides were excluded from the assay. The quantification significance level was as follows: q-value less than 0.05, and absolute log2(fold-change) greater than 0.58 when comparing HD72- MSNs versus C116-MSNs.

### Data Accession

Raw data and complete MS data sets have been uploaded to the Center for Computational Mass Spectrometry, to the MassIVE repository at UCSD, and can be downloaded using the followinglink:https://massive.ucsd.edu/ProteoSAFe/dataset.jsp?task=3a14708986c7468197598 b328d1db750 (MassIVE ID number: MSV000088650; ProteomeXchange ID: PXD030786).

### Western Blot Analysis

C116 and HD72-MSNs were harvested in mammalian protein extracting reagent (150 µl, Thermo Fisher Scientific, 78501) mixed with a protease inhibitor cocktail (1 tablet/10 mL, Roche, 11836170001). Cells were then processed further via sonification using 5 s of pulsing, 5 s of rest for 5 rounds at 40 mA. Samples were then spun down at 14,000 rpm at 4°C for 20 min and quantified using a bicinchoninic acid assay (Thermo Fisher Scientific, 23227). Protein lysates of 10–20 μg were added in with DTT (1 µL) and LDS NuPAGE buffer (6 µL). Proteins were boiled at 95°C for 10 min. Running conditions used were 4–12% Bis-Tris gel in 5% MOPS Running Buffer (Invitrogen, NP0001) at 200 V for 1 h. The transfer conditions used were a 0.45-μm polyvinylidene fluoride (PVDF) membrane transferred in 5% Transfer Buffer (Invitrogen, NP00061) at 20 mA for 840 min. Primary mouse monoclonal antibody Septin-2 (Proteintech, 60075-1-Ig, 1:20,000), rabbit polyclonal antibody Septin-9 (Proteintech, 10769-1-AP, 1:500) and mouse monoclonal IGFBP7 antibody (Santa Cruz Biotechnology, sc-365293,1:100) were incubated at 4°C.

### MSNs Treatment with APOE3 and Lipid Metabolism Quantification

Prepatterned Activin A–treated NSCs from C116 and HD72 were plated at 90,000 cells per well in eight-well chamber slide (Falcon, 354108) for MSNs differentiation using synaptojuice A and B medium. MSNs cultured in serum withdrawal were treated for 48 h with APOE3 (PeproTech, 350- 02-500UG) at 312 ng/ml. Non-treated cells were used as control. For each treatment, a duplicate was performed. The cells were fixed using 4% paraformaldehyde (Sigma, 158127) in 0.1 M PBS, pH 7.4 (Corning, 21-040-CV) for 30 min. To identify the lipid metabolism, the fixed MSNs were stained with Nile Red (Thermo Fisher Scientific, N1142), a lipophilic dye at a dilution of 1/1000 in PBS for 30 min. To quantify the lipid metabolism, a laser scanning confocal microscopy with spectral fingerprinting on a Zeiss LSM 980 was used. The fluorescence emission peak of Nile Red shifts yellow to red, based on increasing polarity of the bound lipid (48). Using automated spectral component extraction, we observed a shorter wavelength punctate (peak at 593 nm) and a longer wavelength (peak at 611 nm) diffuse fluorescence of Nile Red using excitation at 514 nm. Using the “Count cellular foci with secondary cell segmentation“ pipeline in Image Analyst MKII (Image Analyst Software Novato, CA), images taken using a Plan-Apochromat 63×/1.40 Oil lens were segmented using DAPI-stained nuclei as seeds and finding cell boundaries using the diffuse, longer wavelength fluorescence of Nile Red. Then the punctate shorter wavelength foci were counted per cellular area. For analysis of intensities, images recorded with a Plan-Apochromat 20×/0.8 lens and the basic fluorescence histometry using nuclear markers (1–3 labels).

### Pathway Analysis and Network Visualization

Pathway enrichment analysis was performed using g:Profiler with parameters set to *Homo sapiens*, custom background (all proteins identified in FAIMS DDA acquisitions: supplemental Table S2), Benjamini-Hochberg FDR and threshold at 0.05. The gene-sets included for the pathway enrichment analyses were obtained from Gene Ontology (GO) database (GOBP_AllPathways), updated February 01, 2020 (http://download.baderlab.org/EM_ Genesets/). Enrichment results are available in **supplemental Tables S7** and **S8**. The pathway analysis and network visualization were carried out by using Cytoscape 3.7.2 and Cytoscape Enrichment Map application [version 3.2.1 of Enrichment Map software (49) with the following parameters: analysis type = generic/gProfiler, p-value cutoff = 1, FDR Q-value cutoff = 0,05 and similarity between gene-sets was filtered by Jaccard plus overlap combined (coefficient: 0.375)]. The network was manually rearranged to improve layout and clusters of nodes were automatically annotated using the AutoAnnotate Cytoscape App to highlight the prevalent biological functions among a set of related gene-sets.

### Comparison with Multiple Datasets and Drug Prediction

Enrichment analysis for GO biological processes with differentially expressed proteins (FDR < 0.05, logFC > 0.58) was done utilizing the R package clusterProfiler. Drug prediction was done utilizing the LINCS L1000 characteristic direction signatures search engine (https://maayanlab.cloud/L1000CDS2/#/index) [23] with upregulated and downregulated proteins as input (50).

### Experimental Design and Statistical Rationale

Proteomic experiments were conducted with iPSC-derived cultured C116-MSNs (n = 3) and cultured HD72-MSNs (n = 3). “Hyper Reaction Monitoring” retention time peptide standards (HRM, Biognosys, Schlieren, Switzerland; Kit-3003) were spiked into the samples before LC-MS/MS analysis in DDA and DIA modes. More precisely, one DIA cycle (3.2 s) was composed of the acquisition of one MS1 scan, followed by the acquisition of 64 variable windows (5–90 m/z) covering the full mass range (m/z 400–1,250) with an overlap of 1 m/z. DDA and DIA data were used for XIC-based LFQ, performed at the MS1 and MS2 levels, respectively, as detailed hereafter. To determine significantly altered protein groups, pairwise comparisons were performed using a Limma t-test for DDA-based quantification results and a paired t-test for DIA-based quantification, and obtained p-values were corrected for multiple testing, as described hereafter.

## RESULTS

### Generation and Characterization of MSNs from HD-iPSCs

The striatum is dramatically impacted in HD. MSNs, GABAergic inhibitory neurons, are one of the main cell types lost from this region and represent 90% of the striatal neuronal population. To model HD, we used human patient–derived HD-iPSCs (72CAG/19CAG, HD72) that were genetically corrected to a normal repeat length (21CAG/19CAG, C116), thus creating an isogenic control (31). Then, both cell types were differentiated into MSN-like neurons by a method that mimics the major brain developmental stages for this neuronal type: neural induction, regional patterning toward a lateral ganglionic eminence (LGE) identity in presence of Activin A and terminal differentiation (30, 31, 34, 51, 52) (**Fig. 2*A***). Using immunocytochemistry, we found that the cultures were positive for the MSN marker DARPP-32 and neuronal marker MAP2 (**Fig. 2*B***). HD72-MSNs showed less DARPP-32 (p ≤ 0.01) and MAP2 (p ≤ 0.001) than in C116-MSNs (**Fig. 2*C***). This result is consistent with the expression of these markers in postmortem HD striatum and in mouse models of HD (53–55).

**Figure 2.**
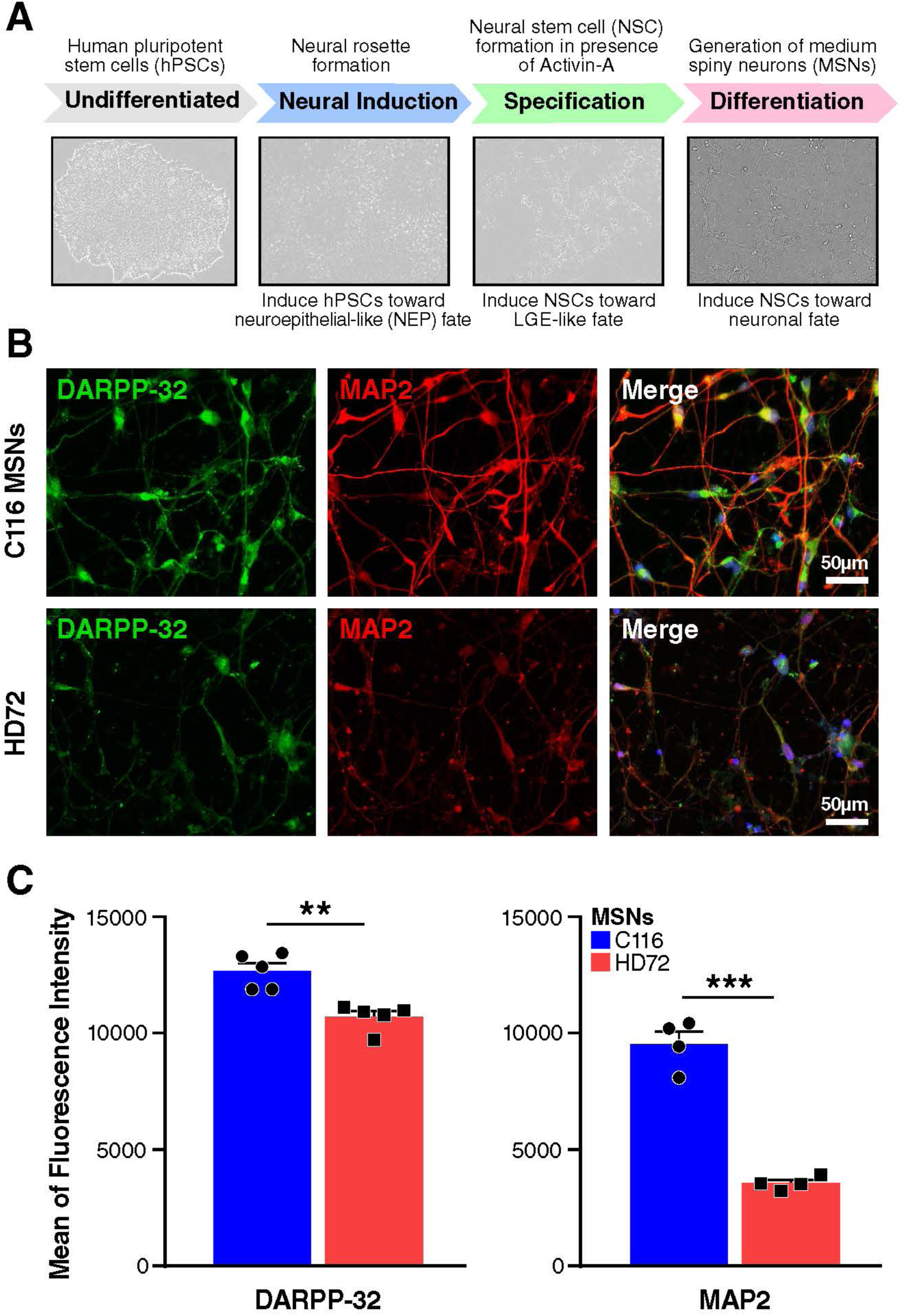
Generation and characterization of iPSC-derived MSNs. *A*, Schematic of steps illustrating the generation of neural stem cells and differentiated MSNs. The method used to differentiate iPSCs into MSNs mimics the major brain developmental stages, including neural induction, regional patterning toward an LGE identity in presence of Activin A, and terminal differentiation. *B*, C116-MSN and HD72-MSN were immunostained after differentiation into MSNs with DARPP-32 (green) and MAP2 (red). Scale bars: 100 µm. *C*, Quantification of the expression levels of DARPP-32 and MAP2 were carried using the Biotek and Image J analysis of the expression levels. Unpaired t-test with Welch’s correction **P ≤ 0.01, ***P ≤ 0.001.

### Proteomic Analysis of HD-MSNs

The HD72- and C116-MSNs were grown in parallel with three replicates for each genotype and subjected to the proteomic workflow in **Fig. 1**. Intracellular proteins were extracted, digested and subjected to a comprehensive quantitative proteomic analysis by LC-MS/MS using a combined approach: protein discovery and LFQ using FAIMS-DDA MS on a Orbitrap Lumos system, followed by protein candidate validation using DIA MS on a TripleTOF 6600 system, and data and bioinformatic analysis (**Fig. 1, supplemental Table S1**).

First, we coupled an additional gas-phase separation using FAIMS ion mobility to DDA acquisitions, and specifically applied three internal CV steps, -50 V, -65 V and -85 V. The gas-phase separation protocol reduces the complexity of the ion population entering the mass spectrometer, which provides deeper MS/MS sampling and proteome coverage (38, 56, 57) (**Fig. 3*A***).This process allowed us to identify 6,323 unique protein groups (≥ 2 unique peptides, FDR ≤ 0.01, **supplemental Table S2**), among which 6,294 protein groups were quantifiable by LFQ algorithms in Proteome Discoverer (**Table 1** and **supplemental Table S3**), providing a comprehensive and deep proteomic dataset for the HD72-MSNs. Assessing the LFQ-MS1-based protein quantification reproducibility within each experimental condition, using three biological replicates of isogenic C116-MSNs and HD72-MSNs, revealed that the coefficient of variation for peptide peak areas was under 20% for 74% of all peptides of the C116-MSN group and 86% of all peptides of the HD72-MSN group (**Fig. 3*B***). Reproducibility of protein group identifications is displayed for three biological replicates of HD72-MSNs (**Fig. 3*C***). FAIMS-DDA quantification details for all replicates are shown in **supplemental Table S3**. Of the 6,294 quantifiable protein groups (using FAIMS-DDA), when comparing HD72-MSNs to C116-MSNs, 901 proteins were significantly changed (**supplemental Table S4*B***): 443 proteins were upregulated, 458 were downregulated (FDR set at 1% and absolute Log2(fold-change) ≥ 0.58), and 5,393 were unchanged (**Fig. 4*A*, Table 1**).

**Figure 3.**
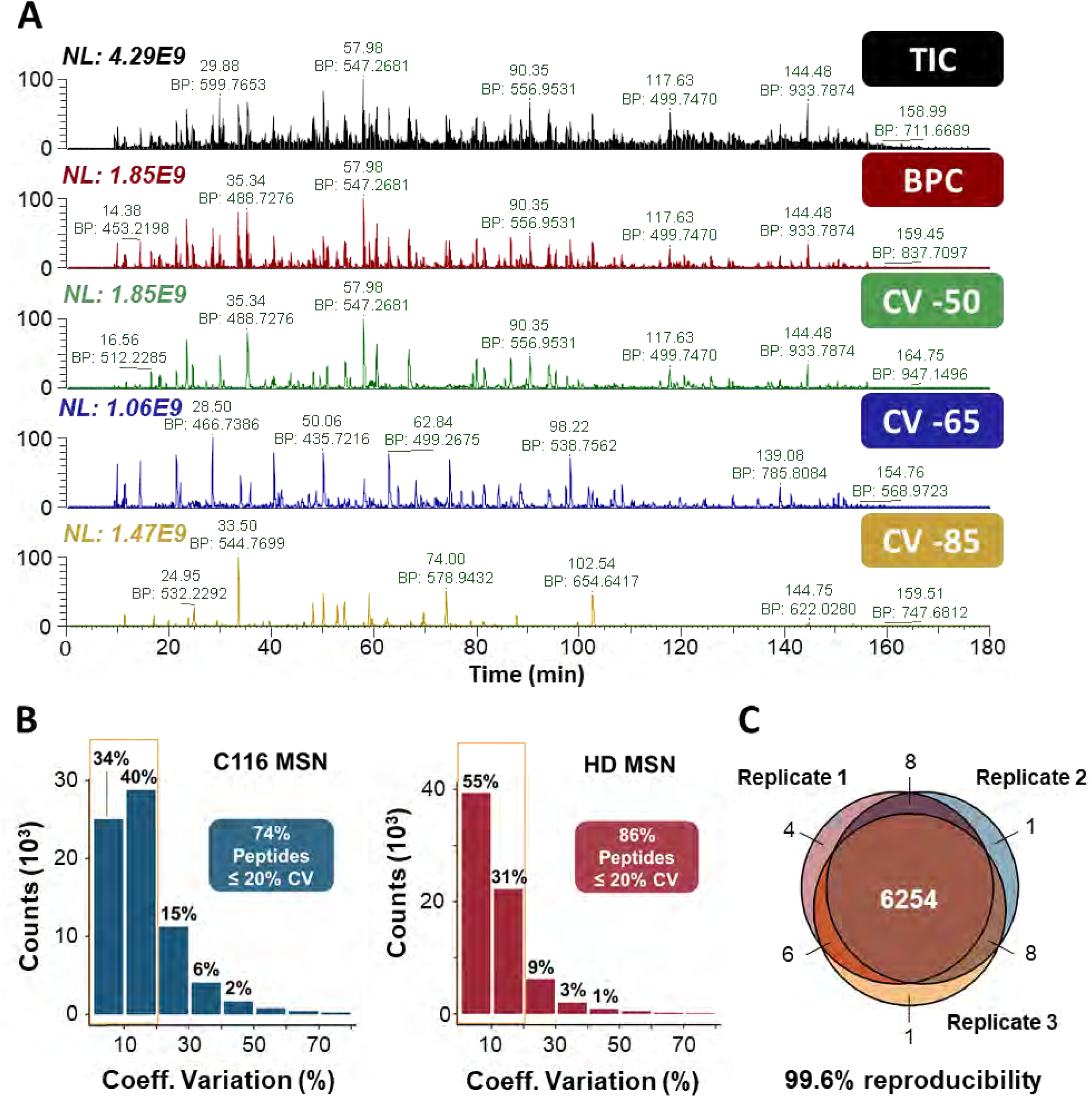
Deep proteome coverage using FAIMS gas-phase separation with DDA: Performance of the FAIMS-DDA MS workflow. *A*, Total ion chromatogram (TIC) and base peak chromatogram (BPC), followed by BPC with the three differential CVs (-50 V, -65 V, and -85 V) for a 2 µg C116-corrected MSN sample injection on a FAIMS-Orbitrap Lumos system operating in DDA mode. *B*, Coefficients of variation (CV) of peptides quantified in three biological replicates of C116-corrected-MSNs and HD-MSNs. *C*, Reproducibility of protein groups identified in three biological replicates of HD-MSNs.

**Figure 4.**
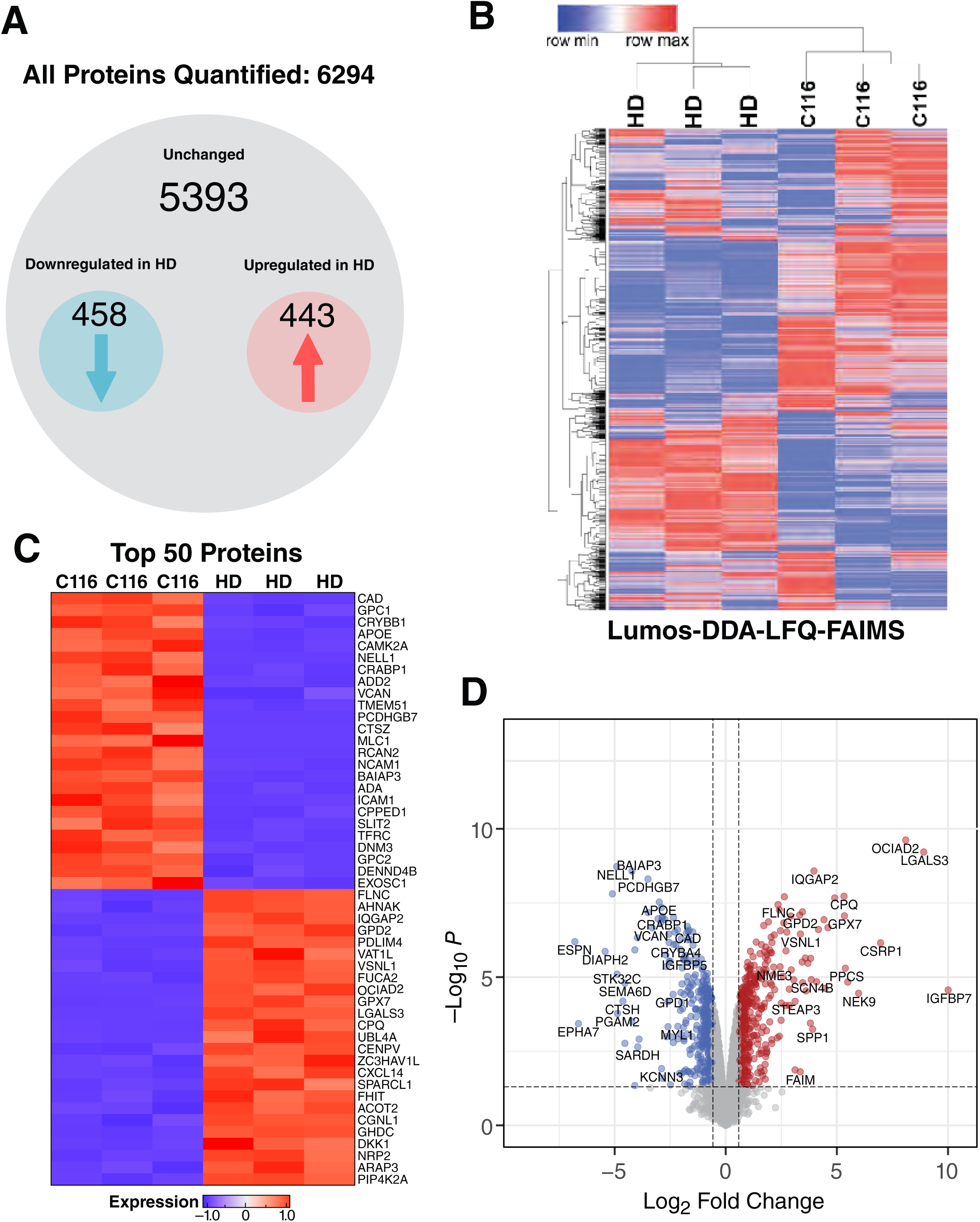
Differential analysis of the proteome of isogenic C116- and HD72-MSNs by FAIMS-DDA MS. *A*, Summary of the proteins quantified and significantly altered using FAIMS gas-phase separation with DDA. *B*, Heat map illustrating the abundance of the proteins of the C116- and HD72-MSNs identified by FAIMS DDA MS with at least two peptides (FDR ≤ 0.01). The heat map represents more precisely the values of MS peak area for n = 3 C116-MSNs and n = 3 HD72-MSNs. *C*, Heat map illustrating the top 50 statistically significant altered proteins in the HD72- versus C116-MSNs using FAIMS-DDA MS. *D*, Volcano plot illustrating the proteins differentially expressed when comparing HD72- versus C116-MSNs (significant proteins: FDR at 1% and log2 fold-change absolute value > 0.58).

To further validate these protein candidates for HD72-MSNs, we used a comprehensive quantitative methodology, DIA-MS, in which fragment ions (MS2) were quantified with accurate relative quantification results. This approach for C116 and HD72-MSN provided protein quantification for 3,106 protein groups with at least two peptides with very high reproducibility (**supplemental Fig. S1**) with at least two peptides identified (**supplemental Table S5**). In fact, 72% and 80% of the identified precursor ions presented a coefficient of variation below 20% for C116-MSNs and HD72-MSNs, respectively (**supplemental Fig. S1**). From the FAIMS DDA discovery study 129 protein candidates were confirmed by the highly quantitative workflow using DIA-MS (q-value ≤ 0.05 and absolute Log2(ratio) ≥ 0.58) (**supplemental Table S4*B*,*C***).

Using hierarchical clustering of protein abundances, we evaluated the variation in C116- MSNs and HD72-MSNs. Heat map representation of the proteomics showed distinct clustering of the two sample groups that depended on CAG length, with HD72-MSN samples being clearly delineated from C116-MSNs (**Fig. 4*B*, supplemental Table S3**). This is consistent with previous studies that found distinct phenotypes for HD and corrected NSCs (32).

To visualize the clustering, quality and significantly altered proteins in HD72-MSNs and C116-MSNs, a heatmap is shown in **Fig. 4*C*** for the top 50 proteins with the highest statistical significance. The biological function, cellular component, molecular function, KEGG, Reactome and WIKI pathways of each protein are summarized in **supplemental Table S2**. The analysis of significantly altered proteins is depicted in the volcano plot showing the estimated Log2(fold-change) versus -Log10(p-value) for each protein, with significantly regulated proteins having a p-value that guarantees a 1% FDR and an absolute Log2(fold-change) value above 0.58 (**Fig. 4*D***, **Table 1**, **supplemental Table S4**). The FAIMS-DDA MS workflow applied for the discovery step resulted in 901 significantly changed protein candidates (**Table 1**), and 129 of these proteins were additionally validated by DIA MS with confidence as significantly changing (**Table 1**).

Newly discovered and previously implicated proteins in HD are shown in the heatmap and volcano plots (**Fig. 4*C***,***D***). One of the top upregulated proteins was insulin-like growth factor-binding protein 7 (IGFBP7). IGFBP7 is released by senescent cells, and cellular senescence is a pathway we previously identified as activated in HD-MSNs with a multitude of relevant markers, including IGFBP7 mRNA (58). IGFBPs are biomarkers for multiple diseases, and their expression causes neurodegeneration (59–65). Western blot analysis further validated the increased levels of IGFBP7 in HD72-MSNs (**supplemental Fig. S2**). OCIAD2, another top upregulated protein, is implicated in Parkinson (PD) and Alzheimer disease (AD) and activates STAT3 (66–68). GPX7 (glutathione peroxidase) is upregulated in HD72-MSNs. GPX activity is increased in HD patient blood (69), and GPX7 (related family member, GPX6) is neuroprotective when overexpressed in HD yeast, *Drosophila* and mouse models (70, 71). The WNT antagonist, Dickkopf1, is also upregulated and is important in striatal synaptic degeneration (72). Proteins involved in lipid metabolism, such as apolipoprotein E (APOE), are downregulated and will be discussed further below (**Fig. 4*D***).

### Functional Enrichment and Protein Network Analysis Reveal Molecular Hallmarks of HD

Functional enrichment studies (49, 73, 74) with the significantly altered proteins in HD72- MSNs revealed molecular dysregulation in a number of pathways (**Fig. 5**). We applied the up- and downregulated protein lists from FAIMS-DDA MS proteomic workflow to g:Profiler with custom background proteins (**supplemental Table S6**) (75). The most significant enriched GO biological process terms upregulated in HD72-MSNs, include ECM-related pathways (e.g., Integrin-Laminin signaling, TGF-beta regulation of ECM, epithelial-mesenchymal transition (EMT) activation, activation of matrix metalloproteinases), cardiovascular system, angiogenesis, Tap63 pathway, DNA replication, senescence, organism development, regulation of cell migration and locomotion, aminoglycan glycosaminoglycan proteoglycan, organism development, regulation of cell migration and locomotion, growth factor stimulus and fatty acid processes **(supplemental Table S7)**. Conversely, processes associated with the downregulated proteins include neurogenesis-axogenesis and, more specifically, the brain-derived neurotrophic factor (BDNF) signaling pathway, Ephrin-A:EphA pathway, regulation of synaptic plasticity, triglyceride homeostasis cholesterol, plasmid lipoprotein particle immune response, INF-γ signaling, immune system MHC complex, triglyceride homeostasis, lipid metabolism, lymphocyte proliferation and cellular response to stimulus (**Fig. 5**). Pathways involved in organism development and regulation of cell migration and locomotion, and regulation of biological and homeostatic processes were both up- and downregulated. Although HD is predominantly associated with dysregulation of the CNS, HD affects the peripheral tissues, which results in symptoms, such as cardiomyopathy, skeletal muscle malfunction, weight loss, metabolic and immune dysregulation (76, 77). Our proteomic analysis demonstrated that cardiomyopathy, cardiovascular and angiogenesis related pathways are upregulated in HD72-MSN (**Fig. 5**, **supplemental Table S7**).

**Figure 5.**
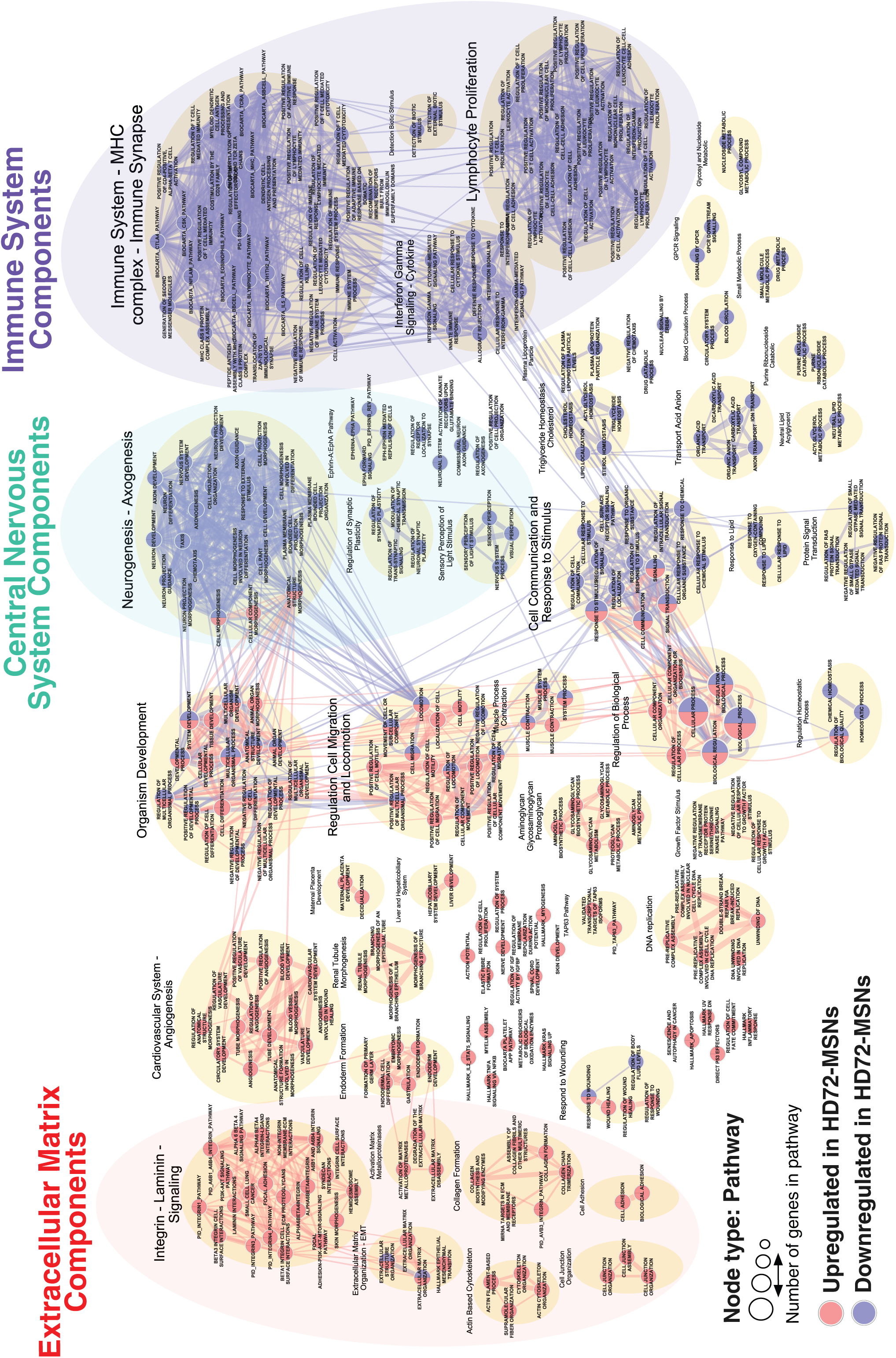
Functional enrichment map of significantly altered proteins in HD72-MSNs. Enrichment results from g:Profiler were mapped as a network of gene sets (nodes) related by mutual overlap (edges) in Cytoscape using its EnrichmentMap. Red color represents gene sets upregulated in HD72-MSNs, whereas blue represents gene sets downregulated in HD72-MSNs. Node size is proportional to the gene-set size, and edge thickness represents the number of overlapping genes between sets. Functional enrichment map was generated with proteins quantified by FAIMS-DDA MS.

The canonical pathways from a complementary analysis using Ingenuity Pathway Analysis (IPA) are shown in **supplemental Fig. S3.** The top pathways include hepatic fibrosis, semaphorin neuronal repulsive signaling pathway, regulation of cellular mechanics of calpain protease, IL-4 signaling, axonal guidance signaling, caveolar-mediated endocytosis signaling, SNARE signaling, estrogen receptor, RHO GTPase signaling, CLEAR signaling and many more that have been implicated in HD.

### Reactome Functional Interaction Network for Isogenic HD-MSNs Upregulated Proteins

Next, we used functional interaction analysis to define clusters of proteins that are closely connected to each other with ReactomeFIViz, a reactome functional interaction network (78). We identified an interconnected network with 142 of the 443 upregulated proteins (**Fig. 6**, **supplemental Table S8*A***). There were 11 clusters for the upregulated proteins with extracellular matrix and DNA signaling (DNA replication pathway, double-stand break repair, G1/S transition) having the highest significance (**Fig. 6*A,B***). We describe each cluster below, along with its correlation with functional enrichment and the relevant HD pathogenic mechanisms.

**Figure 6.**
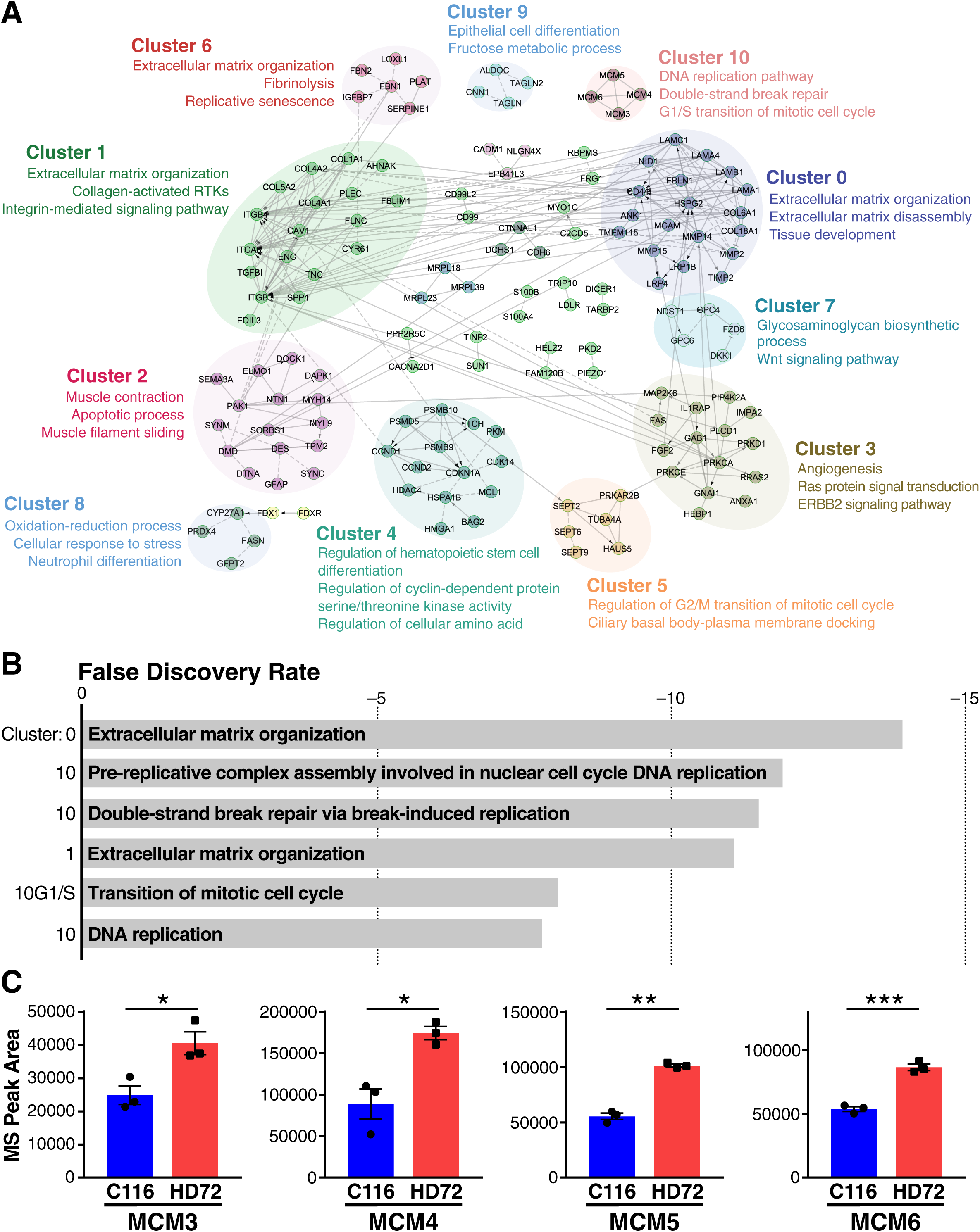
*De novo* sub-network construction and clustering using proteins differentially upregulated when comparing HD72-MSNs to C116-MSNs. *A*, Networks of genes were constructed using 443 upregulated proteins in HD72-MSNs as determined using FAIMS-DDA MS. The functional network and clustering were performed using the Reactome Functional Interaction Network (ReactomeFIViz). Nodes in the network correspond to genes, and edges correspond to interactions. Shaded ovals represent clusters of genes sharing common enriched biological functions. *B,* Classification of clusters based on false discovery rates. *C*, Quantitative proteomics reveals MCM3, MCM4, MCM6 and MCM6 are expressed more highly in HD72-MSNs than C116- MSNs. Unpaired t-test with Welch’s correction *P ≤ 0.05, **P ≤ 0.01, ***P ≤ 0.001.

### Clusters 0, 1, 4, 6 *–* Extracellular Matrix Organization, Replicative Senescence

The upregulated interaction network confirmed the dysregulation of ECM-regulated pathways in HD72-MSNs. The Cluster 0 protein network was associated with ECM organization, disassembly and tissue development, highlighting the role of matrix metalloproteinases in HD72- MSNs: CD44, MMP2, MMP14 and TIMP2 (**Fig. 6*A***, **supplemental Table S8*A***). Cluster 1 network identified key proteins involved in ECM-organization, collagen and integrin signaling: ITGA6, ITGB4, ITGB3, COL1A1, COL4A1, COL4A2, COL5A2, CAV1, FLN and TNC (**Fig. 6*A***). ECM-related pathways represent a large group comprising ECM organization, integrins, laminins, collagens, cell adhesion and hemidesmosome signaling. The development of CNS requires neuronal proliferation, axon guidance and synaptic communication to ensure the proper formation of neuronal networks. These processes are highly coordinated by the ECM ligands and their specific cell adhesion receptors (79). Therefore, dysregulation of ECM components impairs the formation and maintenance of neural circuitry and increases risk for several neurological pathologies, such as HD, AD, and autism spectrum disorder (80). We previously described the role of MMP14 and TIMPs in HD (81, 82).

Normal HTT has a role in the construction and regulation of the ECM, and its absence results in disruption of ECM components (83). Our results support the hypothesis that the abnormal polyQ expansion within the mHTT affects the ECM components that ensure the integrity of MSNs in terms of neuronal identity, architecture, and ability to interact with neighboring cells.

This hypothesis is consistent with the detection of EMT pathways in the upregulated proteins (**Fig. 5**, **supplemental Table S7**). EMT is a critical cellular process in embryonic development that enables epithelial cells to acquire the properties of mesenchymal cells. ECM proteins are important in maintaining epithelial integrity, in addition to initiating and regulating the EMT (84). HD is characterized by impairment of specification and maturation of MSNs (85). mHTT may impair age-dependent maintenance of striatal MSN identity gene expression (86). Our proteomic analysis showed a dysregulation of ECM-related pathways with the increased EMT-related proteins, which may indicate the inability of HD72-MSNs to acquire and maintain a neuronal signature. Notably, we found that CNS-related pathways were downregulated in HD72-MSNs, including neurogenesis, axonogenesis, axon guidance, regulation of axonal synapse activity, dopamine, and glutamine metabolic process (**Fig. 5**, **supplemental Table S7**).

In clusters 4 and 6, we also identified proteins involved in cellular senescence, including CDKN1A (p16), SERPINE1 and IGFBP7. Recent studies in mouse models of AD and PD suggest cellular senescence is important in disease progression and pathogenesis (10, 87–91).

### Clusters 2, 3 *–* Muscle Contraction, Netrin Signaling and Angiogenesis

Interestingly, clusters 2 and 3, Netrin/SEMA1 signaling, Ras protein, Erbb2 signaling, and angiogenesis were novel dysregulated pathways in HD72-MSN that were not identified in pathway enrichment analysis (**Fig. 6*A***). IPA analysis also identified these pathways, and the networks are shown in **Supplemental Fig. 3**.

### Cluster 5 *–* Septin Signaling Pathways in HD

Notably, three members of the septin protein family, SEPT2, SEPT6 and SEPT9, were present in cluster 5. Septin proteins participate in various physiological processes, such as cytoskeleton regulation, cell division, membrane trafficking, neuronal formation, and maintenance. Septin dysregulation is associated with diverse diseases, including cancer, infection, and neurological disorders (92, 93). Using quantitative proteomics, we found that SEPT2, SEPT3, SEPT4, SEPT5 and SEPT9 are dysregulated in HD MSN (**Fig. 7*A***). Further, western blot analysis on HD72- and C116-MSNs provide validation of the proteomic results. Both SEPT2 and SEPT9 were upregulated in HD72, compared to C116-MSNs (**Fig. 7*B* and *C***, p ≤ 0.05, p ≤ 0.01).

**Figure 7.**
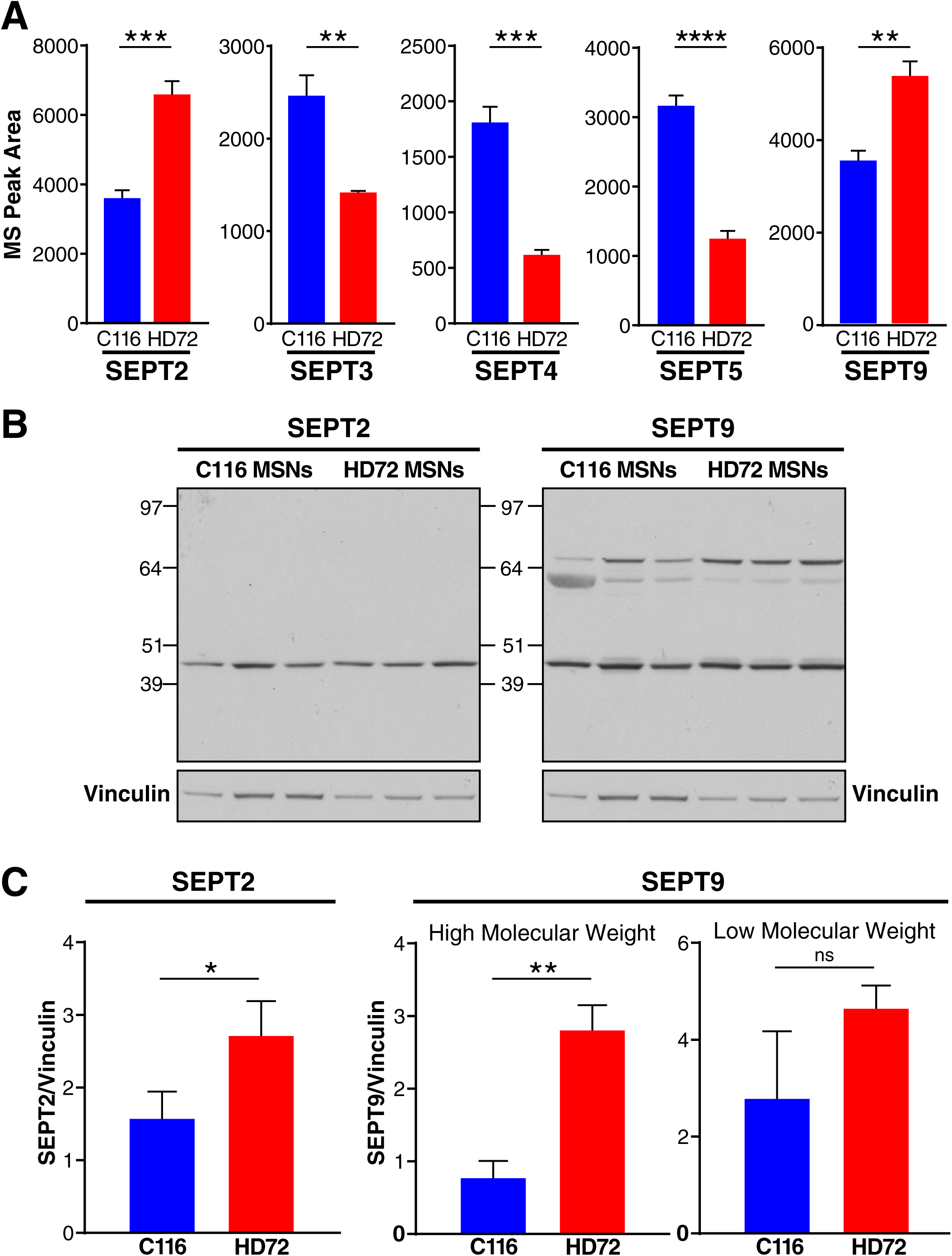
SEPTIN family members are dysregulated in HD-MSNs. *A*, Quantitative proteomics reveals SEPT2, SEPT3, SEPT4, SEPT5 and SEPT9 are dysregulated in HD-MSNs. *B*, Western blot analysis shows that SEPT2 and SEPT9 were upregulated in HD72-MSNs, compared to C116-MSNs. *C*, Quantification of SEPT2 and SEPT9 levels, normalized to vinculin. Unpaired t- test with Welch’s correction *P ≤ 0.05, **P ≤ 0.01, ***P ≤ 0.001, ****P ≤ 0.0001, ns, not significant.

### Cluster 7 *–* Glycosaminoglycan Biosynthetic Process and Wnt Signaling

Cluster 7 identifies glycosaminoglycan biosynthetic process and Wnt signaling. The Wnt signaling is altered in human and mouse models of HD and may be an early event in the pathogenesis of HD (94–96). The Wnt signaling pathway is also enriched in the downregulated proteins. Interestingly, glycosaminoglycan biosynthetic process is dysregulated in AD. A major components of the ECM are glycosaminoglycans. These include heparan sulfates and chondroitin sulfates. The growth factors (e.g., FGF, BDNF, HB-EGF) bind glycosaminoglycan regulating their activity. The ECM, particularly, the sulfated glycosaminoglycan component, is structurally and functionally altered in AD hippocampus(97).

### Clusters 8,9 *–* Oxidation Reduction Process and Fructose Metabolic Process

Clusters 8 and 9 highlight that proteins associated with fatty acid oxidation and fructose metabolic processes were largely upregulated in the HD-MSNs. Fatty acid metabolism is dysregulated in HD and linked to reduction of active sterol regulatory element responsive protein 2 (SREBP-2) (98–100). Fructose metabolism is linked to levels of uric acid, and levels of uric acid in biofluids are lower in HD patients than controls (101).

### Cluster 10 *–* DNA Signaling Is a Top Enriched Pathway in HD MSNs and Implicates MCM Proteins

Cluster 10 is pre-replicative complex assembly involved in nuclear cell-cycle DNA replication. Genome-wide association studies suggest genes involved in DNA-damage-repair mechanisms are modifiers of the age of onset and disease severity in HD. Further, HTT acts as a stress-response protein to modulate DNA damage. We identified the minichromosome maintenance (MCM) proteins 2–7 as a top dysregulated pathway. The MCM complex regulates DNA replication, cell-cycle and DNA damage responses, and so, it is likely an important signaling pathway for HD (102–104). Cluster 10 showed MCM3, 4, 5 and 6 as dysregulated pathways in HD-MSNs (**Fig. 6*A,B***). Analysis of individual MS peaks confirmed that levels of MCM3, 4, 5 and 6 are significantly greater in HD-MSNs than C116-MSNs (**Fig. 6*C***).

### Reactome Functional Interaction Network for Isogenic HD-MSNs Down-Regulated Proteins

We identified an interconnected network with 147 of the 458 down-regulated proteins (FAIMS-DDA MS) (**Fig. 7**, **supplemental Table S8*B***) using ReactomeFIViz (78). The down- regulated related clusters were associated with immune system-related pathways, axon guidance signaling, MAPK cascade, calcium modulation and the Wnt signaling pathway (**Fig. 8*A***). Correspondingly, for downregulated pathways, antigen processing and presentation, interferon-gamma signaling and ephrin receptor signaling were the most significant (**Fig. 8*B***). We describe each cluster below, including its correlation with functional enrichment and relevance to HD pathogenic mechanisms.

**Figure 8.**
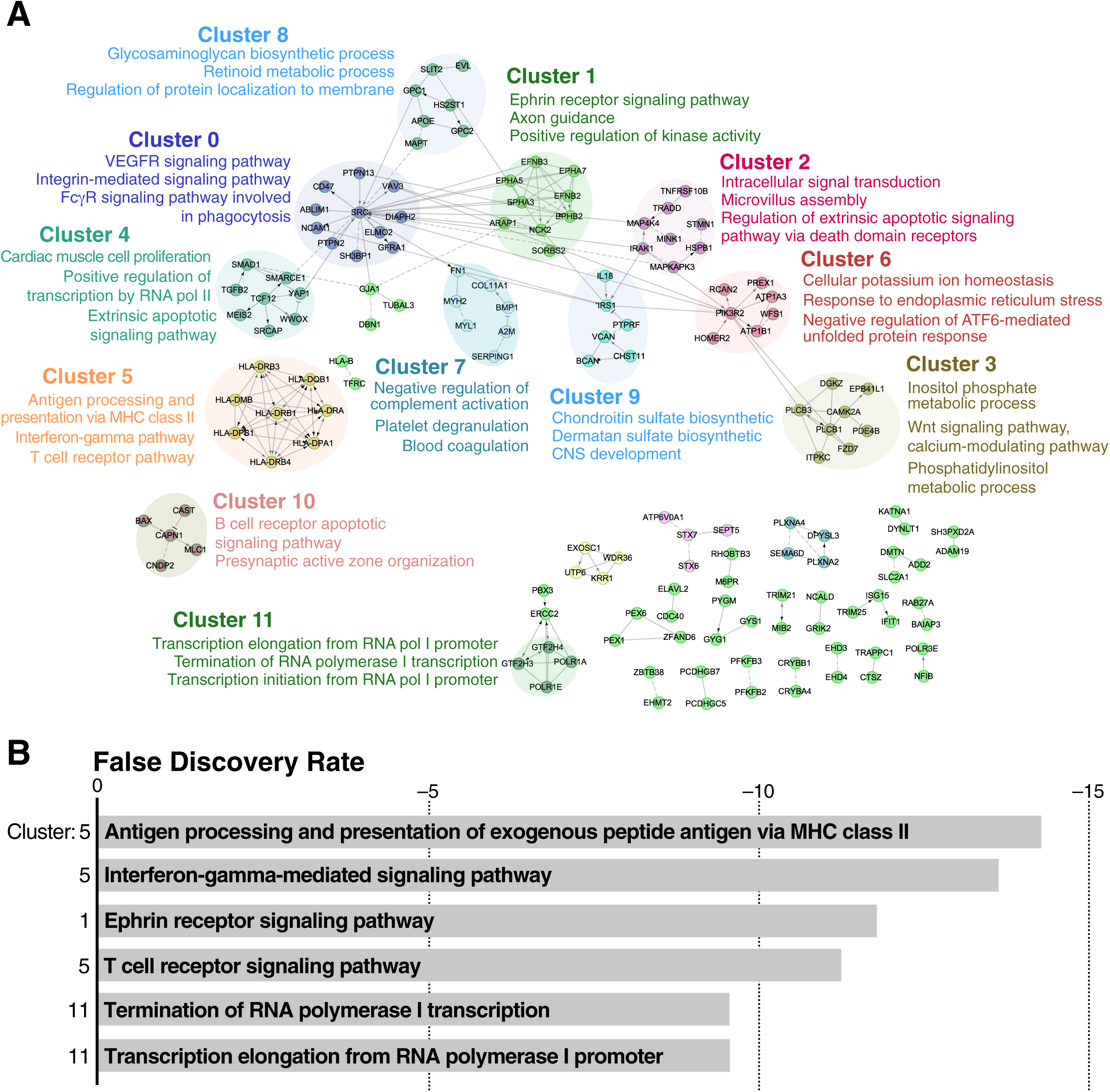
*De novo* sub-network construction and clustering using proteins differentially downregulated when comparing HD72-MSNs to C116-MSNs. *A*, Network of gene was constructed using 458 downregulated genes in HD72-MSNs as determined using FAIMS-DDA MS. The functional network and clustering were performed using the Reactome Functional Interaction Network (ReactomeFIViz). Nodes in the network correspond to genes, and edges correspond to interactions. Shaded ovals represent clusters of genes sharing common enriched biological functions. *B,* Classification of clusters based on false discovery rate.

### Clusters 0 *–* VEGFR, Integrin and FcγR Signaling Pathways

Alterations in VEGFR and VEGR are common in a number of triplet repeat diseases and modulation of VEGR is neuroprotective in HD. Depletion of this growth factor may contribute the HD-MSN phenotype. The Fc gamma receptors (FcγRs) are generally thought to be expressed in immune cells but are also expressed in neurons where they may mediate excitatory pathways, and therefore are downregulated in HD-MSNs.

### Clusters 1, 2 *–* Axonal Guidance through Ephrin Signaling and the Stathmin Pathway

Previous transcriptomic studies of HD-derived NSCs identified altered pathways related to neuronal development, axonogenesis and axonal guidance (32, 33). Our proteomic analysis of HD-MSNs is consistent with the hypothesis that mHTT prevents the proper development and maintenance of MSNs by downregulating processes related to CNS development (**Fig. 5**) (86). Furthermore, the functional interactions analysis identified the downregulation of axonal guidance pathway by impairing ephrin proteins, including EPHA5, EPHA7, EPHB2 and EPHA3 (**Fig. 8*A and B)***. We also found that stathmin-1 (STMN1), a protein belonging to the stathmin family, is downregulated in HD-MSNs (**Fig. 8*A*, Cluster 2**). STMN1 is mainly expressed in the CNS, including the striatum, with high expression during the neuronal development, maturation and plasticity (105, 106). Dysregulation of STMN1 occurs in neurological disorders, including AD (107), amyotrophic lateral sclerosis (108) and spinal muscular atrophy (SMA) (109). AAV9 gene therapy that overexpressed STMN1 in an SMA mouse model led to a reduction in the SMA phenotype including increased survival and weight (109).

### Clusters 3, 8 *–* Dysregulation of APOE Signaling and Lipid Metabolism in HD72-MSNs

Pathway enrichment analysis revealed a dysregulation of lipid metabolism, including triglyceride and cholesterol pathways in the HD72-MSNs, when compared to the corrected C116- MSNs (**Fig. 5**). Correspondingly, APOE expression was downregulated in HD-MSNs and likely modulates the lipid metabolism alterations in HD72-MSNs (**Fig. 8*A***). Further, cluster 3 links HD to altered inositol phosphate metabolism and phosphatidylinositol metabolic processes. Numerous studies link HD to alterations in lipid metabolism, and IPA analysis suggested APOE is a top upstream regulator (**supplemental Fig. 3**). In addition, several of the lipid metabolism proteins significantly altered in HD-MSNs are regulators of lipid droplet formation (110). These include monoacylglycerol lipase, lipid droplet–associated hydrolase, diacylglycerol O-acyltransferase, low density lipoprotein receptor, NPC intracellular cholesterol transporter 2 and sodium-coupled neutral amino acid transporter. To quantify for lipid-rich components, including lipid droplets, we used a lipophilic dye Nile Red and discriminated for neutral lipid and phospholipid (**Fig. 9*A,B***). In serum-withdrawal culture, we found that HD72-MSNs had significantly higher level of lipid droplets (**Fig. 9*C***), and a trend towards increase in neutral lipids and phospholipids (**supplemental Fig. 4**), compared to C116-MSNs. Treatment with APOE3 increased the numbers of lipid droplets in HD72 more than in C116-MSNs (**Fig. 9*B,C***). This suggests HD72-MSNs respond and modify lipid metabolism distinctly from control cells. In addition, we showed that HD72-MSNs have lower levels of APOE than C116-MSNs (**Fig. 9D,E**). These data indicate that HD72-MSNs have dysregulated levels of endogenous APOE, in association with the ability to accumulate a significant number of lipid droplets.

**Figure 9.**
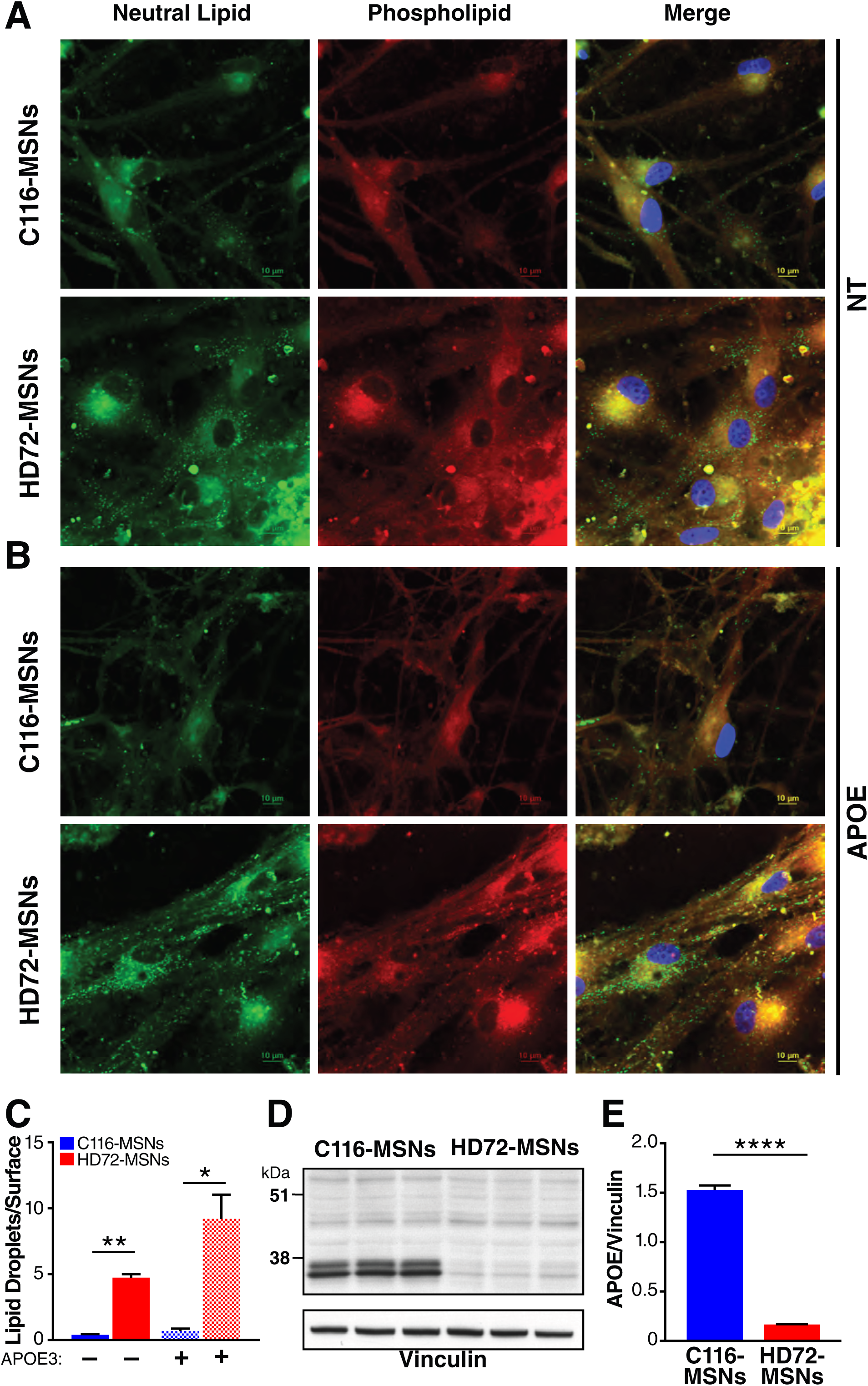
HD-MSN lipid metabolism and its modulation by APOE3. C116-MSNs and HD72- MSNs cultured in serum withdrawal were treated for 48 h with 312 ng/mL APOE3. Non-treated (NT) cells were used as controls. The cells were stained with Nile Red, a lipophilic dye. *A,B*, Representative confocal images for lipid droplets. *C*, Quantification of the density of the lipid droplets in normal and APOE3 treated condition. Scale bar: 10 µm. *D*, Western blot analysis of APOE in C116-MSNs and HD72-MSNs. *E,* Quantification of APOE levels normalized to vinculin. Unpaired t-test with Welch’s correction *P ≤ 0.05, **P ≤ 0.01, ****P ≤ 0.0001.

### Cluster 4 – TGFβ, SMAD and TCF12 Signaling Pathways

Cluster 4 identifies TGFβ2 and SMAD1 proteins as deregulated in HD72-MSNs (**Fig. 8*A***), consistent with previous studies in HD-NSCs (32, 33). Our results do not address how TGFβ and SMAD activities affect the temporal acquisition of the striatal fate but suggest this may be relevant differentiation into MSNs (111). Interestingly, cluster 4 also contains TCF12 protein, a member of the basic helix-loop-helix protein family that regulates early and late events of neuronal differentiation (112, 113). TCF12 interacts with another proneuronal basic helix-loop-helix protein, NEUROD1, and thus contributes to the migration as well as maturation and survival of newborn neurons (114). HTT interacts with NEUROD1 and a small molecule Isx-9 that upregulates NEUROD1, correcting aberrant signaling and guidance in HD-NSCs (33). Our results suggest that TCF12 is also a potential target in HD-MSNs (**Fig. 8*A***).

### Clusters 5, 7 *–* Downregulation of HLA- and CNS-Related Proteins in HD-MSNs

Intriguingly, pathway enrichment and functional interaction analysis illustrated that CNS-related pathways are downregulated in HD72-MSNs along with the immune system and IFN-γ responses (**Fig. 5**, **8**). Although the CNS was considered to be immunologically inert, the major histocompatibility complex class I (MHC-I) is expressed in mouse brain and neurons (115–118). Recently, using a single-cell transcriptomic, Darmanis et al. showed that MHC-I genes are expressed in a subset of neurons in the human adult brain (119). Therefore, our results show the expression of MHC components and the IFN-γ response are dysregulated in HD72-MSNs. The functional interactions analysis of downregulated proteins within cluster 0 identified key proteins involved in these pathways: HLA-DPA1, HLA-DPB1, HLA-DMB, HLA-DRA, HLA-DRB1, HLA- DRB4, HLA-DRB3 and HLA-DQB1 (**Fig. 8, supplemental Table S8*B***). Immune system and IFN- γ responses were downregulated in HD72-MSNs, and this raises the intriguing possibility that neurons use this innate immunity signaling system to define neuronal differentiation. Interestingly, MHC-II and DARPP-32 levels were lower in HD72- than C116-MSNs (**supplemental Fig. S5**).

### Clusters 6, 9 – Sodium and Potassium Ion Homeostasis and Proteoglycan Pathways

Cluster 6 was enriched with proteins related to sodium and potassium ion transport across the plasma membrane, including ATP1A3 and ATP1B1. ATP1A3 is highly expressed in the CNS and critical during brain development by modulating osmotic equilibrium and membrane potential (120). Dysregulation of ATP1A3 is associated with polymicrogyria, a developmental malformation of the cerebral cortex (120). Further investigation of the role of those ionic pumps may shed the light on electrophysiological functions in HD.

Interestingly, in cluster 9, we found that chondroitin sulfate proteins, such as VCAN and BCAN, are downregulated in HD-MSNs. Those proteins may illustrate the function of proteoglycan during important processes of neurodevelopment, such as neuronal migration, differentiation and maturation (121).

### Cluster 11 *–* Transcriptional Initiation, Elongation, and Termination from the RNA Polymerase 1 Promoter

Altered RNA polymerase I activity is consistent with studies showing ribosomal transcription is regulated by PGC-1 alpha, and this process is impaired in HD (122). HTT is also part of a transcription-coupled DNA complex formed by RNA polymerase II subunit A, basic transcription factors, PNKP, ATXN3, DNA ligase 3, CREB protein (CBP, histone acetyltransferase), and this complex identifies lesions in the template DNA strand and mediates their repair during transcriptional elongation. mHTT likely disrupts RNA polymerase I activity in a similar complex (123).

### Comparison of HD-MSNs Proteomic Data Set to Human and Mouse HD Proteome and Modifier Data Sets

We compared our proteomics data set to published proteomics of postmortem HD cortex and knock-in HD mouse model Q175 striatum (**Fig. 10*A***) (24, 86). Enrichment analysis of downregulated proteins showed dysregulation in all three data sets for regulation of small GTPase-mediated signal transduction, regulation of neurotransmitter levels, regulation of neuronal synaptic plasticity, regulation of GTPase activity, regulation of cell morphogenesis involved in differentiation, regulation of axonogenesis, neurotransmitter transport, neuron projection guidance, negative regulation of neuron projection development, negative regulation of cell projection organization, axonogenesis, and axon guidance. Enrichment analysis of upregulated proteins showed alterations in regulation of GTPase-mediated signal transduction, regulation of cell morphogenesis and axonogenesis. Comparison of differentially expressed proteins in the postmortem cortex of HD patients (24) and those in MSN showed an overlap of 50 proteins. In contrast, comparison the Q175 mouse proteome (86), a well-established HD model, and the striatum had an overlap of 19 proteins. This suggests there is a closer correlation at the protein level between human MSN and HD postmortem tissue. Analysis with Enrichr indicates our data set overlaps with transcriptomics of HD grade 3 caudate nucleus GSE3790 with a p-value of 2.0e-27 with 147 genes in common.

**Figure 10.**
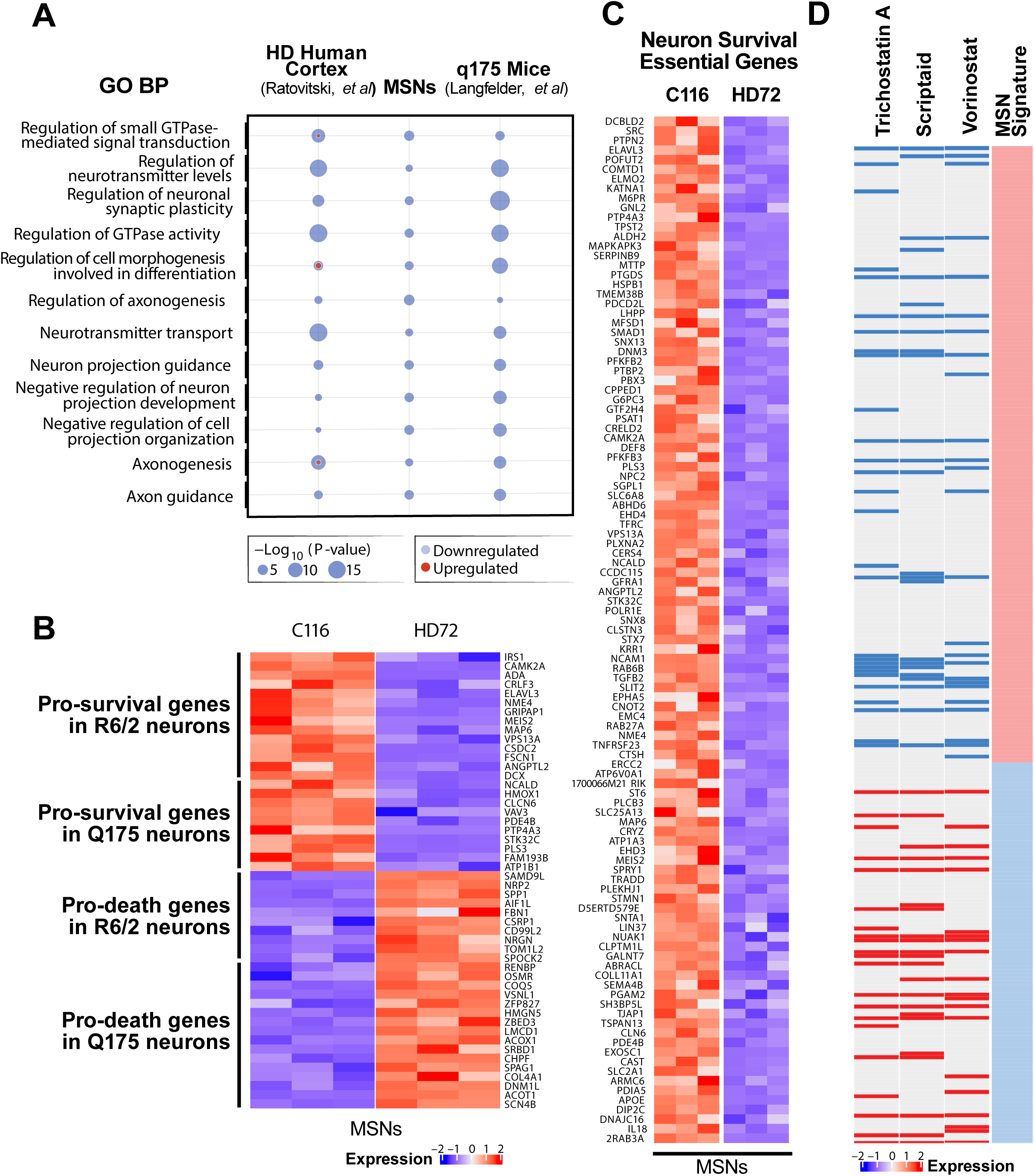
Comparison of HD proteomics data sets. *A*, Comparison of dysregulated GO Biological Process (BP) terms among MSNs from human cortex of HD patients (24) and the Q175 mouse model of HD (86). *B*, Heatmap displaying expression levels of genetic modifiers of HD (133). Pro-death genes in Q175 and R6/2 were compared to upregulated genes in HD72-MSNs, and pro-survival genes were compared to downregulated genes in HD72-MSNs. *C*, Heatmap displaying gene expression levels of downregulated genes in HD72-MSNs to genes found to be essential for neuron survival (133). *D*, The signature of differentially expressed proteins in MSNs was used as input on the L1000CDS2 (124) website to identify drugs that would modify the proteomic signature from HD to corrected. The top drugs identified are shown and are known modifiers of HD. The heatmap on the top shows the proteomic signature of HD and how it is predicted to be affected by each of the drugs.

We also compared the overlap of HD MSN proteomics data set with a published shRNA in vivo screen for modifiers of neuron survival in R6/2 and zQ175 mice (**Fig. 10*B* and *C***). Many known pro-survival proteins in R6/2 or zQ175 mice are down-regulated in our HD72-MSN proteome. These include CAMK2A, ADA, ELAVL3 and GRIPAP2. Correspondingly, we also found pro-death genes upregulated in the HD72-MSN proteome. Particularly striking was the number of neuronal essential genes that are downregulated in HD72-MSN proteome (**Fig. 10*C***). We also compared our HD72-MSN proteome with known HTT protein interactors. This revealed over 50 known HTT protein interactors that are altered in the HD MSN proteome (**supplemental Fig. S6**). Hub proteins that are highly enriched in our proteomic data set includes YWHAB, GRIN2B, ITGB1, APP and DLG4; these are proteins implicated in the pathogenesis of HD.

### Drug Signature of HD-MSNs

We utilized the L1000CDS2 website to identify possible drugs that are predicted to reverse the proteomic signature of HD72-MSNs into C116-MSNs **(Fig. 10*D*)** (124). Trichostatin A, Scriptaid and Vorinostat were among the top hits identified. They belong to classes of histone deacetylase inhibitors that which are beneficial in HD models (125, 126). An extensive list of drugs identified can be found in **supplemental Table S9**.

## DISCUSSION

Using quantitative proteomic analysis by LC-MS/MS with FAIMS for protein discovery (DDA-MS) we provide a comprehensive coverage of the MSN proteome with 6,294 quantifiable proteins identified. Of these proteins, we found ∼14% of the identified proteins had altered expression in HD72-MSNs, compared to isogenic control C116-MSNs. HTT has numerous cellular functions, and the polyQ expansion in the protein would be expected to disrupt multiple pathways involved in neuronal homeostasis. Because these studies were carried out on human neurons derived from HD patient iPSCs, the data may prove useful for further understanding the biology and therapeutic targets in HD MSNs. The comparison of our proteomics data set to published proteomics of postmortem HD cortex showed a strong correlation for down regulated proteins identifying small GTPase-mediated signal transduction, regulation of neurotransmitter levels, regulation of neuronal synaptic plasticity, regulation of GTPase activity, regulation of cell morphogenesis involved in differentiation, regulation of axonogenesis, neurotransmitter transport, neuron projection guidance, negative regulation of neuron projection development, negative regulation of cell projection organization, axonogenesis, and axon guidance.

Our data set has a robust enrichment for processes involved in the brain function including neurogenesis-axogenesis, the BDNF-signaling pathway, Ephrin-A:EphA pathway, regulation of synaptic plasticity, axonal guidance signaling, caveolar-mediated endocytosis signaling, SNARE signaling and RHO GTPase signaling. The septin family members were particularly interesting as the polyQ expansion in HTT dysregulated multiple family members SEPT2, 3, 4, 5 and 9. Septin family members form highly organized pre and postsynaptic supramolecular structures and regulate synaptic transmission. Misregulation of human septins has been linked to AD and PD (127). SEPT5 is a substrate for ubiquitin ligase Parkin, and the PD loss of function mutations in Parkin lead to the accumulation of SEPT5 and dopamine-dependent neurodegeneration. Correspondingly, SEPT4 knockout mice exhibit reduced dopaminergic neurotransmission. Interestingly, SEPT9 modulates cargo entry into dendrites by regulating the motility of two distinct kinesin motors (128). In HD MSNs, we found SEPT2 and SEPT9 had increased expression, whereas SEPT3, 4, and 5 were downregulated in HD MSNs. The supramolecular structure formed at the synapse for this family of proteins is likely disrupted in HD MSNs. The IPA pathway of DNA replication, recombination, repair and DNA metabolism for the HD72-MSN proteome is shown in **Supplemental Fig. 7**.

One of the top statistically significantly enriched pathways was DNA signaling (DNA replication pathway, double-stand break repair, G1/S transition). This fits with numerous studies in the field suggesting an increase in DNA damage in HD, defects in DNA repair mechanisms and a multitude of genes that enhance or prevent CAG expansion modifying the age of onset of HD (129, 130). We identified the MCM proteins 2–7 as a top dysregulated pathway. The MCM complex regulates DNA replication, cell-cycle and DNA damage responses, and so, it is likely an important signaling pathway for HD (102–104).

One of the signaling pathways identified in our analysis is cellular senescence. We previously demonstrated senescence features develop in human HD NSCs and MSNs (58). p16^INK4a^ promotes cellular senescence in these human HD cells. FOXO3, a major cell survival factor that represses cell senescence, opposing p16^INK4a^ expression via the FOXO3 repression of the transcriptional modulator ETS2. In our current study, we find the genes CDKN1A (p21), IGFBP7, HMGA1 and SERPINE1 are part of the cellular senescence pathway activated in HD72- MSNs. Senescent cells also have a senescence-associated secretory phenotype (SASP). Interestingly, when we compare out data set to the recently defined “SASP Atlas”, a data base of the secretomes of senescent cells (131), we found the following SASP proteins elevated in HD-MSNs: FLNC, AHNAK, CD44, HSPA1B, HSPA1A, TNC, TIMP2, TKT, EMILIN1, COL6A1, HSPG2, TPM2, TAGLN2, PLEC, MMP2, PKM, HMGA1, ALDOC, CALD1, PSAP, YWHAE, and MIF.

Guided by pathway analysis of the HD-72 MSNs proteomics identifying lipid metabolism and genes involved in lipid droplet formation, we found an increase in lipid droplets in HD72- MSNs. The role of lipid droplets in neurological diseases and the brain are not completely understood (132). Our results indicate that HD72-MSNs have dysregulated levels of endogenous APOE, in association with the ability to accumulate a significant number of lipid droplets. Lipid droplets accumulate during aging, inflammation, oxidative stress and in neurological diseases (ALS, AD, PD). This newly discovered phenotype for HD neurons can be utilized in drug and CRISPR screens to find modifiers of this interesting pathway.

Our quantitative unbiased proteomics analysis of HD-MSNs provides a comprehensive understanding the proteins altered in human HD-MSNs derived from patient iPSCs. We identified signaling pathways not previously known to be dysregulated in proteomic of human HD neurons, including MHC class proteins, IFN-γ, cellular senescence, ApoE signaling/lipid metabolism and regulation of cellular response to heat in neurons. Since many of the proteins discovered in our study have been identified in related neurological diseases, our findings will likely accelerate the identification of new biomarkers for HD.

## Acknowledgments

This work was supported by the National Institutes of Health awards: K99 AG065484 (PI: Basisty), S10 OD028654 (PI: Schilling), and 1S10 OD016281 (Buck Institute and R01-NS100529 (PI : Ellerby). Support was also provided by “The Taube Family Program in Regenerative Medicine Genome Editing for Huntington’s Disease” to LME. BS would like to thank Thermo Scientific for scientific and technological support and expertise. KTT was supported by a fellowship from the Collaborative Center for X-linked Dystonia Parkinsonism (CCXDP).

## Data Availability

Raw data and complete MS data sets have been uploaded to the Center for Computational Mass Spectrometry, to the MassIVE repository at UCSD, and can be downloaded using the following link: https://massive.ucsd.edu/ProteoSAFe/dataset.jsp?task=3a14708986c7468197598b328d1d <u>b750</u> (MassIVE ID number: MSV000088650; ProteomeXchange ID: PXD030786).

## TABLES

Table 1. The number of proteins identified using each mass spectrometer (Discovery and Quantification Validation).

**Supplemental** Table S1. DIA window isolation scheme of the DIA MS acquisition method.

**Supplemental** Table S2. Protein, peptide and peptide spectrum match (PSM) identification results obtained from the FAIMS-DDA MS data set.

**Supplemental** Table S3. Protein quantification and statistical analysis results obtained from the FAIMS-DDA MS data set.

**Supplemental** Table S4. Comparison and validation of the significantly changing protein candidates obtained from the FAIMS-DDA MS data set with the significantly changing proteins obtained from the DIA MS data set.

**Supplemental** Table S5. Protein quantification and statistical analysis results obtained from the DIA MS data set.

**Supplemental** Table S6. Custom background of proteins.

**Supplemental** Table S7. Pathway enrichment analysis of the upregulated and downregulated proteins from the FAIMS-DDA MS proteomic analysis.

**Supplemental Table S8A**. Reactome functional interaction network analysis of the upregulated proteins in HD72-MSN to define clusters of proteins that are closely connected.

**Supplemental Table S8B**. Reactome functional interaction network analysis of the downregulated proteins in HD72-MSN to define clusters of proteins that are closely connected.

**Supplemental** Table S9. Predicted drugs that modify HD signature.

## FIGURE LEGENDS

**Supplemental Fig. 1. Performances of the TripleTOF 6600 DIA MS workflow.** DIA data were processed using the pan-human spectral library (45). *A,* Coefficients of variation (CV) of precursors identified in three biological replicates of C116-corrected MSN and HD-MSN. 72% and 80% of the identified precursor ions presented a coefficient of variation below 20% in C116- corrected MSNs and HD-MSNs, respectively. 42% and 54% of the identified precursor ions presented a coefficient of variation below 10% in C116-corrected MSN and HD MSN, respectively. *B*, Number of missing values obtained for each precursor ion in the dataset. 15,531 of a total of 31,183 precursor ions were identified and quantified in all six samples of the dataset (complete profile).

**Supplemental Fig. 2. Insulin-like growth factor-binding protein 7 levels are increased in HD72-MSNs.** *A*, Western blot analysis of IGFBP7 in C116-MSNs and HD72-MSNs. *B*, Quantification of IGFBP7 protein levels normalized to vinculin. Unpaired t-test with Welch’s correction *P ≤ 0.05.

**Supplemental Fig. 3. IPA analysis of the differentially expressed proteins in HD72-MSNs compared to controls.**

**Supplemental Fig. 4**. **HD-MSN lipid metabolism and its modulation by APOE3.** C116- and HD72-MSNs cultured in serum withdrawal were treated for 48 h with 312 ng/mL APOE3. Non-treated cells were used as controls. The cells were stained with Nile Red, a lipophilic dye. *A,B*, Quantifications of the phospholipid (red) and neutral lipid (green) intensity in normal and APOE3 treated condition, as illustrated from the **Fig. 9**. Unpaired t-test with Welch’s correction *P ≤ 0.05, **P ≤ 0.01, ***P ≤ 0.001, ns, not significant.

**Supplemental Fig. 5. Modulation of MHC and IFN-γ rescues HD cellular phenotypes.** C116- MSNs and HD72-MSNs were treated for 48 h with IFN-γ at different concentrations. Non-treated cells were used as control. Representative immunofluorescence images showing that the expressions of DARPP-32 and MHC-II (*A-B*) were modulated after IFN-γ treatment. *C,D*, Quantifications of the expression levels of each marker based on the analysis of pixel intensity were carried using the Biotek and Image J. Scale bar: 200 µm. One-way ANOVA for multiple comparisons (Tukey’s) ****P<0.0001.

**Supplemental Fig. 6. HD protein expression overlaps with HTT interacting proteins.** The overlapping network between significantly altered proteins from FAIMS-DDA MS and the HTT protein interactome identified with *A*, yeast two-hybrid method (134) or *B*, Weighted Correlation Network Analysis (135) or *C*, *in vitro* affinity pull-down assays using mouse protein (7) or *D, in vitro* affinity pull-down assays using human protein (7). *E*, The overlapping network between significantly altered proteins as determined by DIA MS and HTT proteomic interactome identified with Weighted Correlation Network Analysis (135).

## Notes

### Competing Interest Statement

The authors have declared no competing interest.

