## Supplemental Figure 1-7 for "Proteomic Analysis of Huntington’s Disease Medium Spiny Neurons Identifies Alterations in Lipid Droplets"

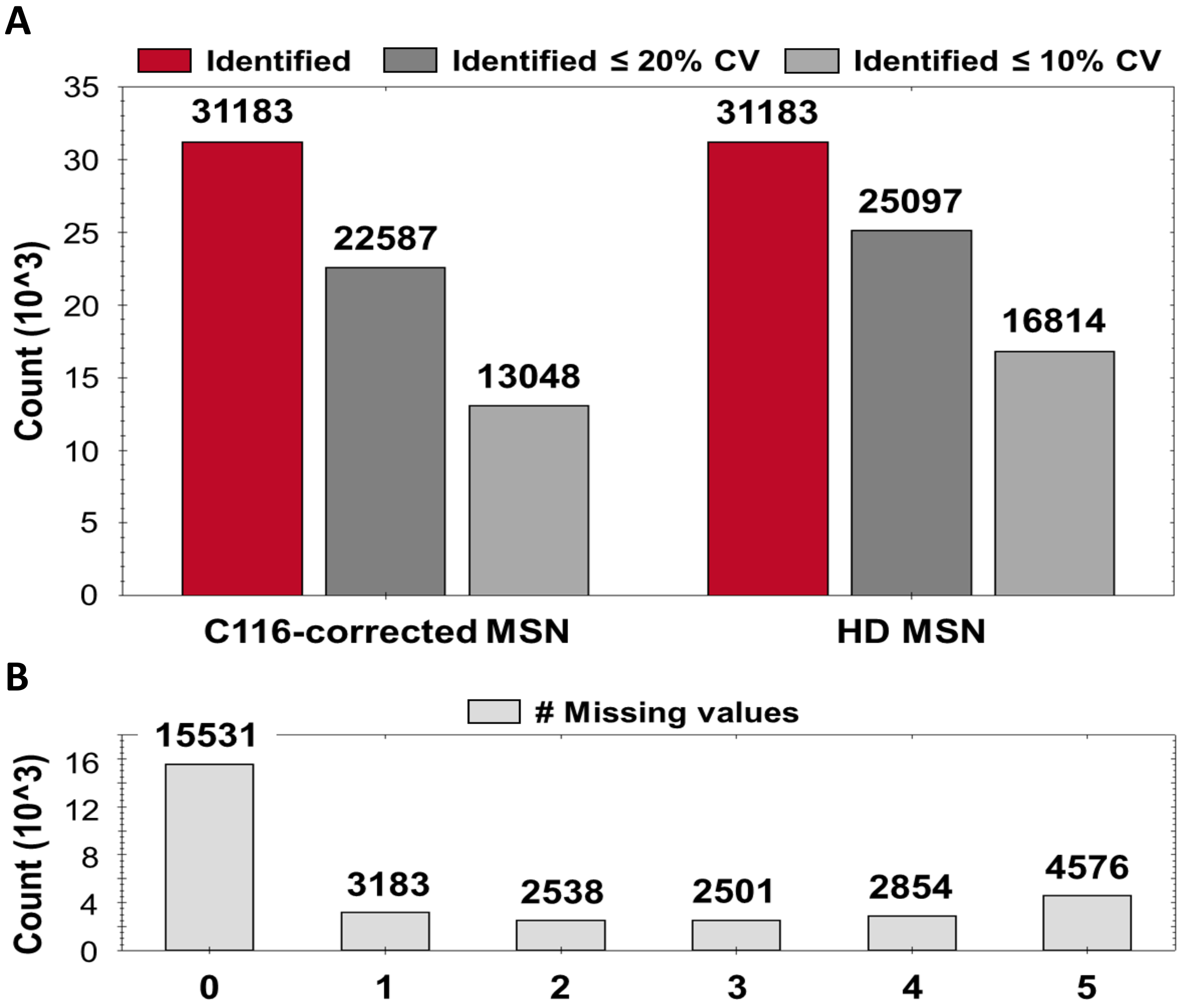


Supplemental figure 1

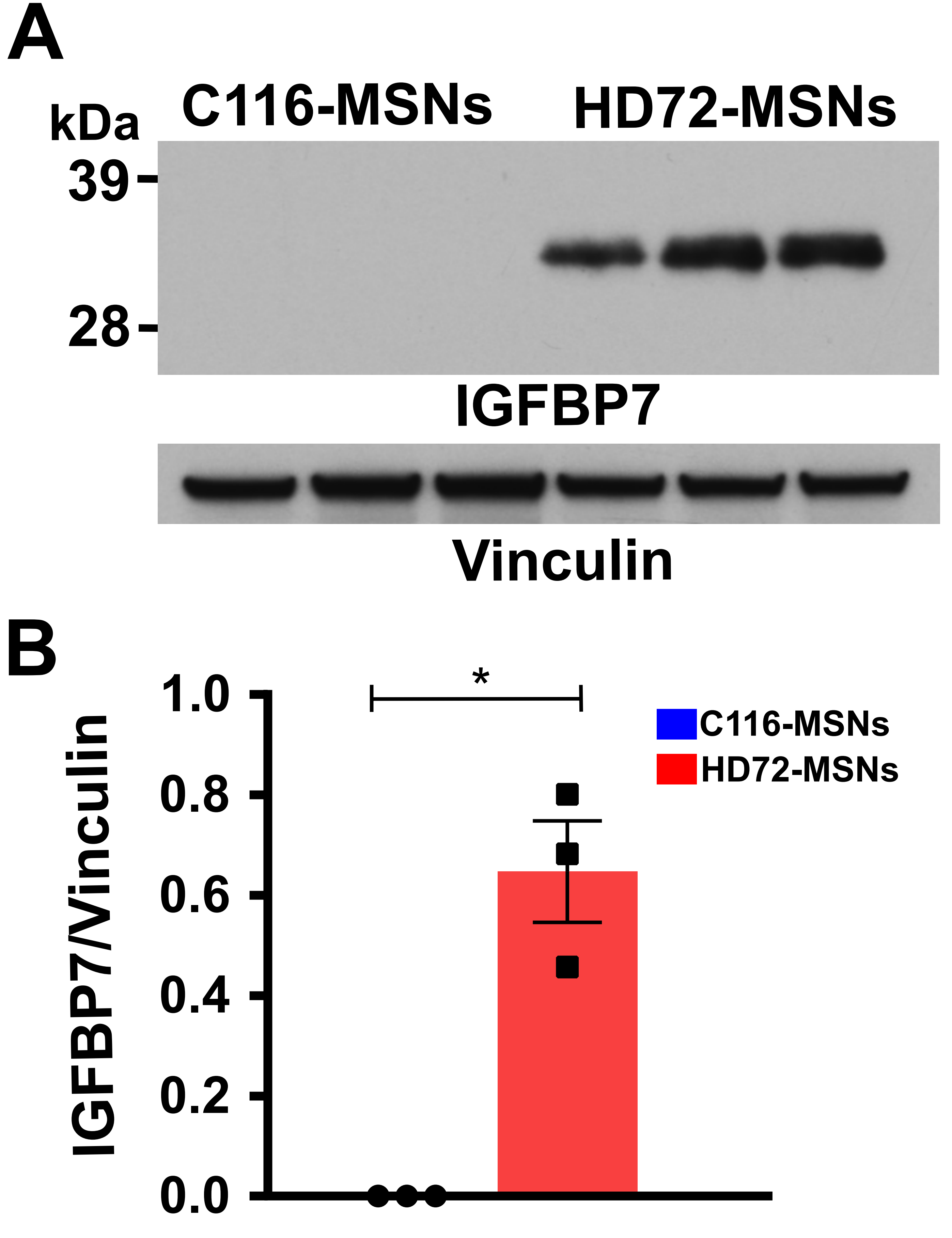


Supplemental figure 2

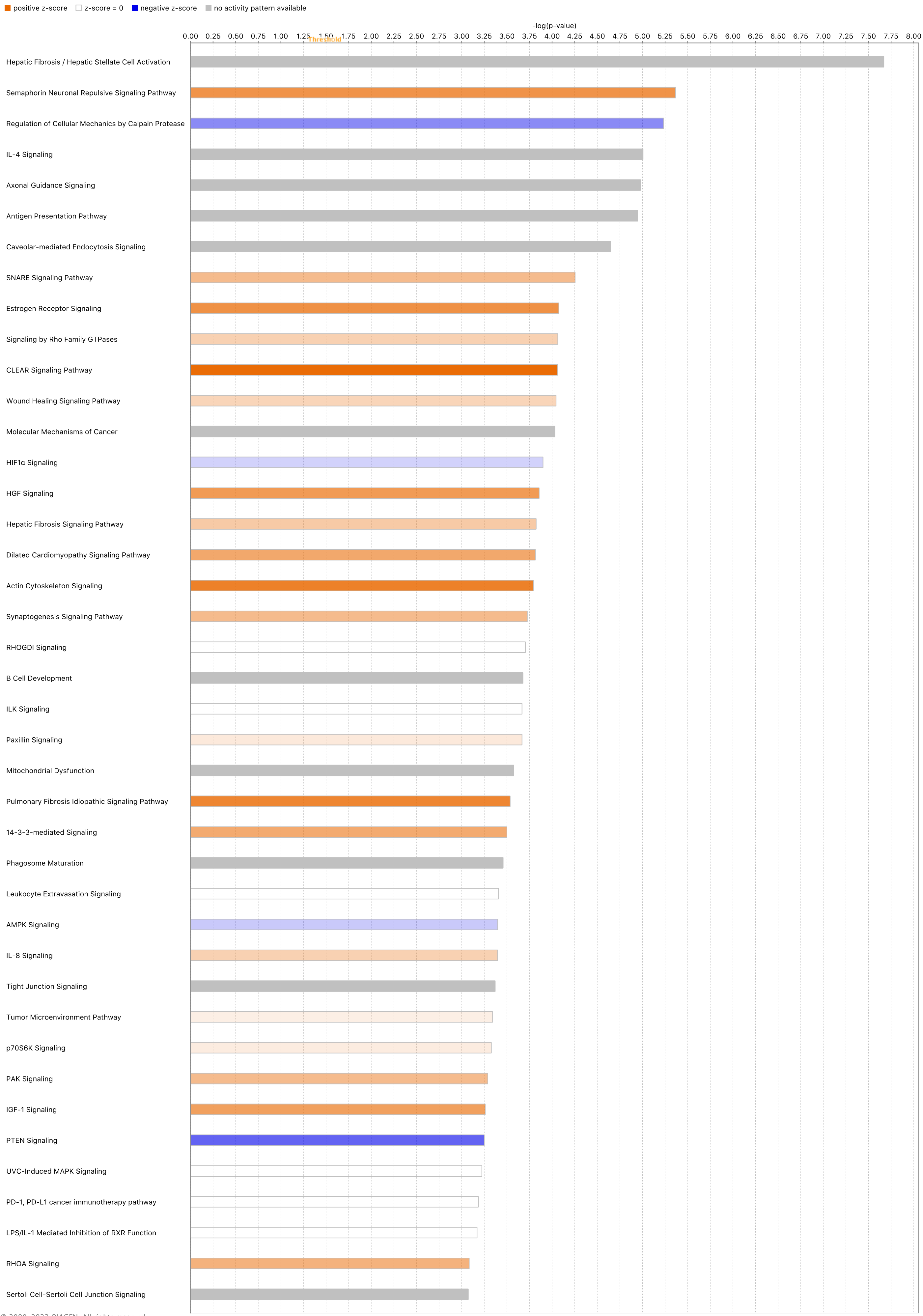


© 2000-2022 QIAGEN. All rights reserved.

**Supplemental Figure 3**

A

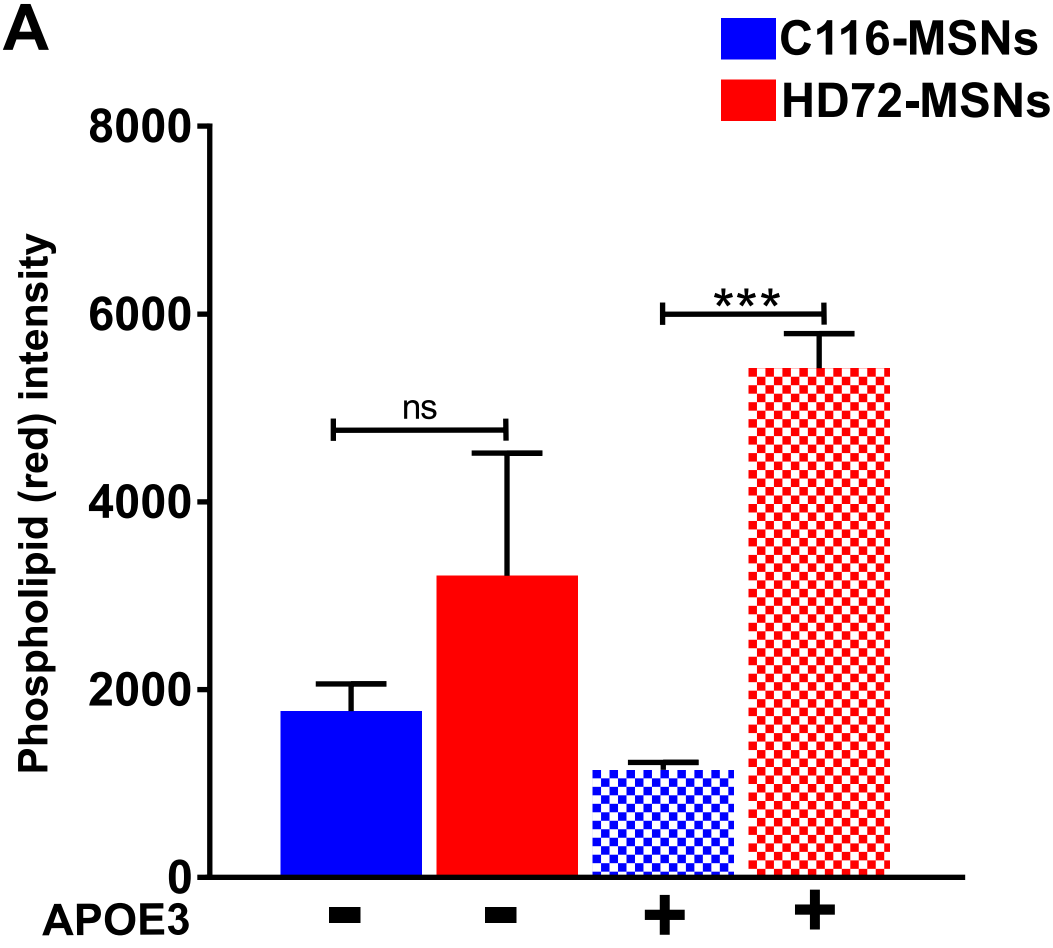


B

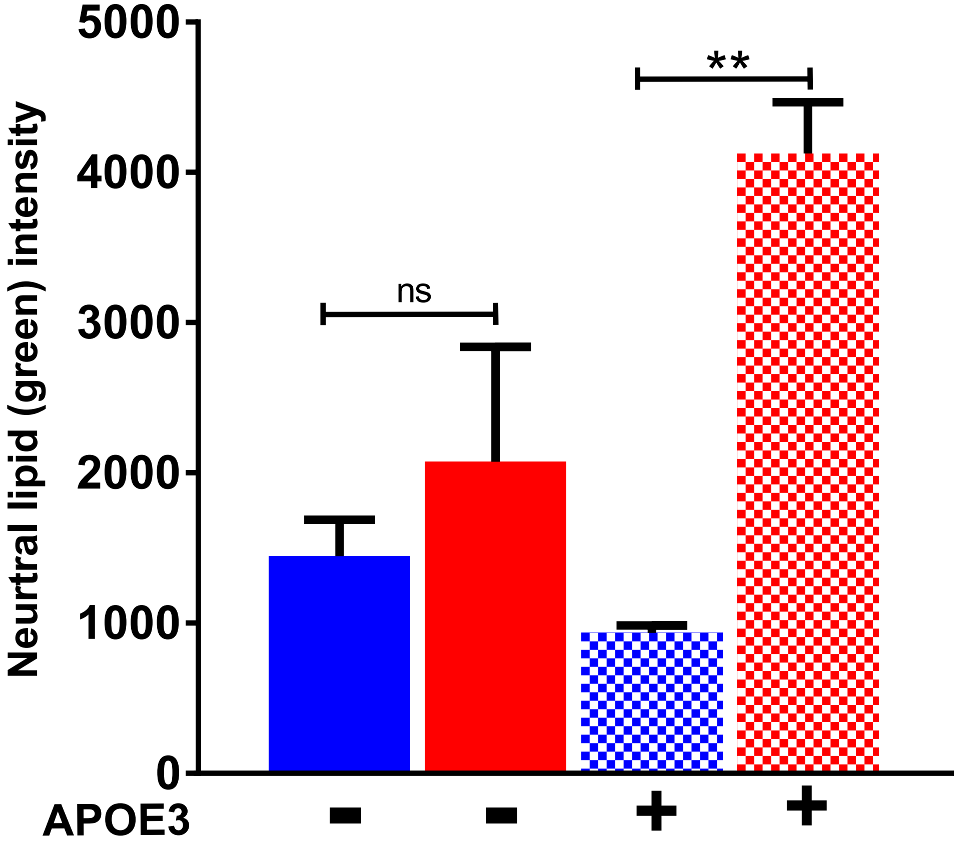


Supplementary figure 4

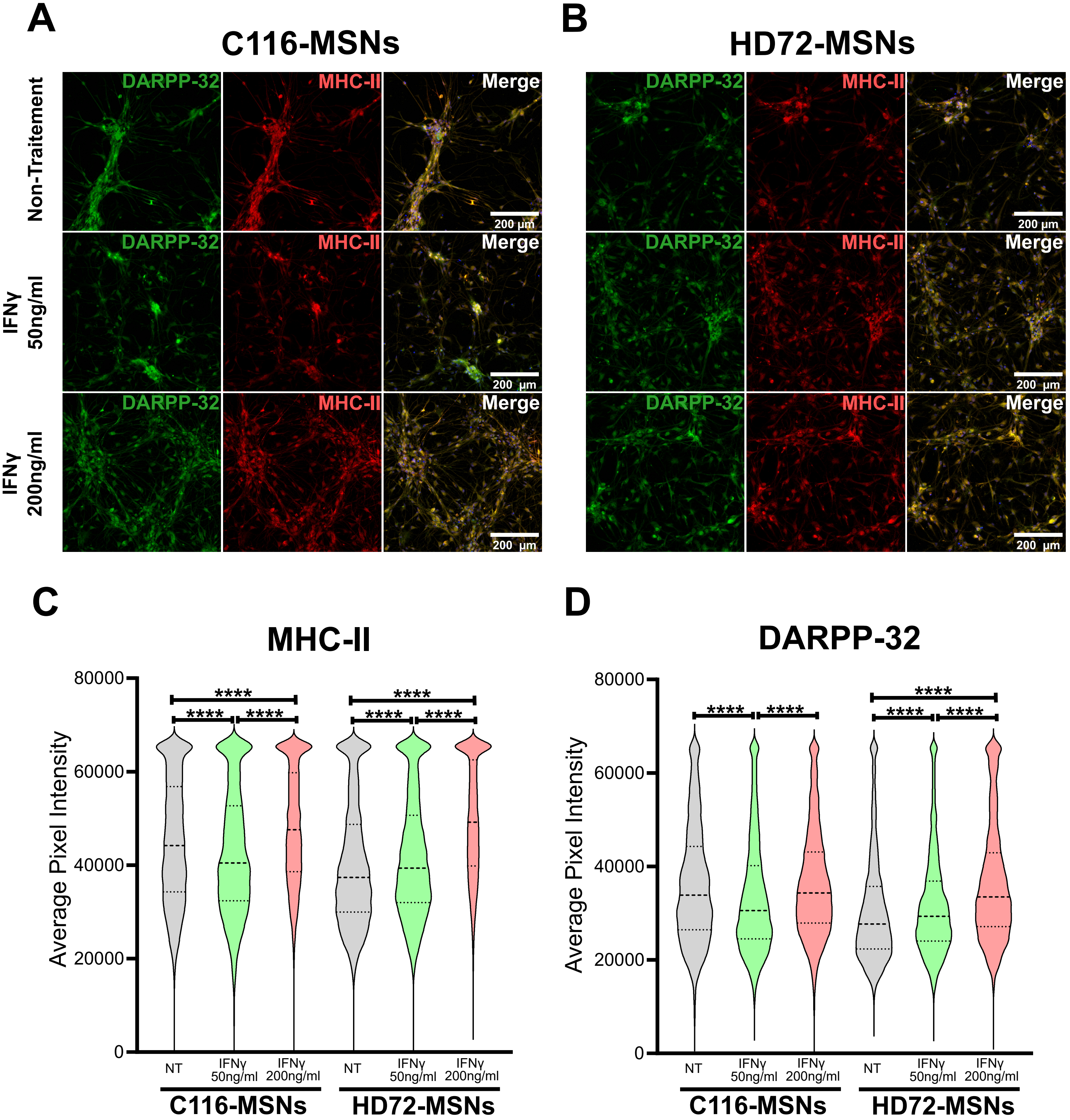


Supplementary figure 5

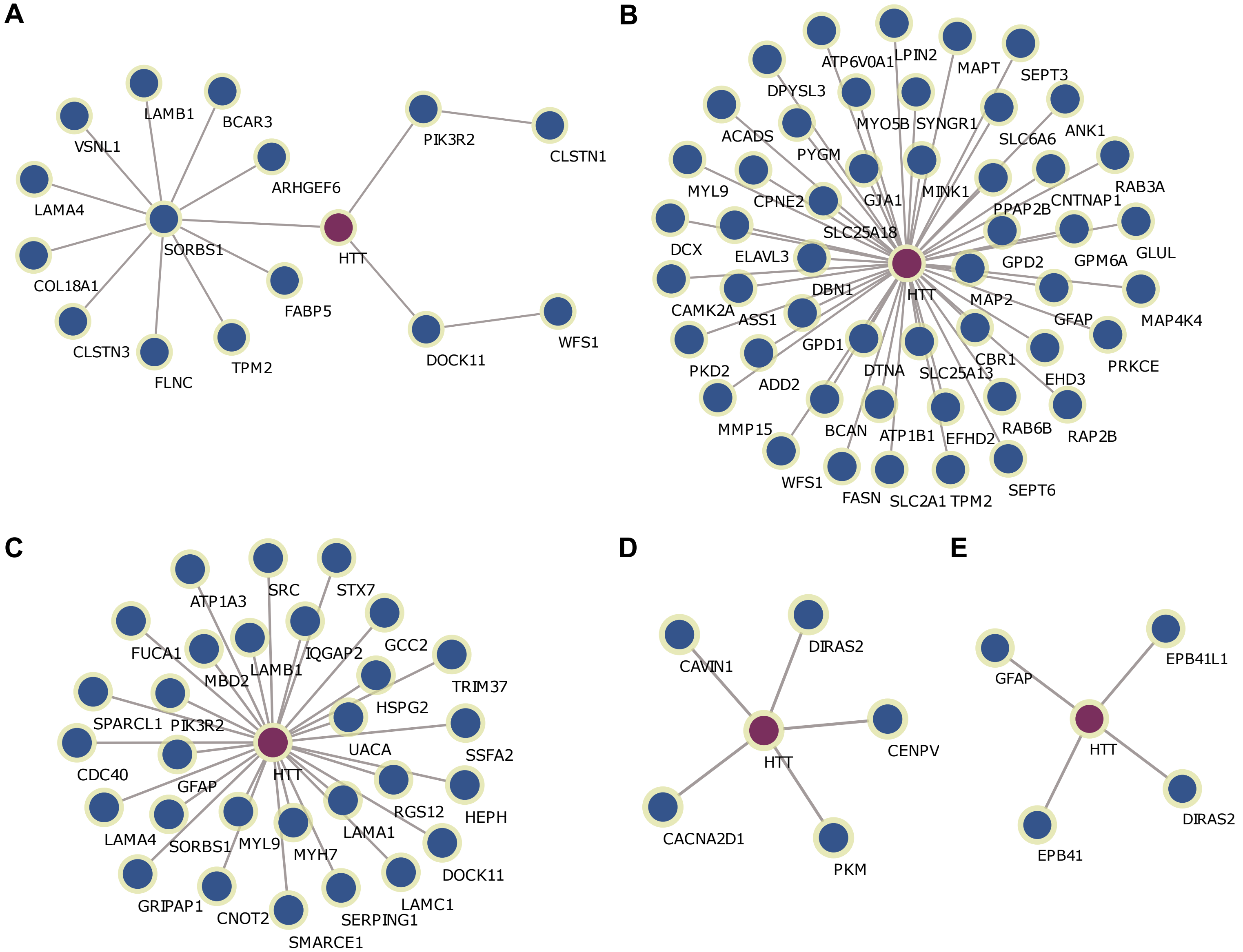


Supplementary figure 6

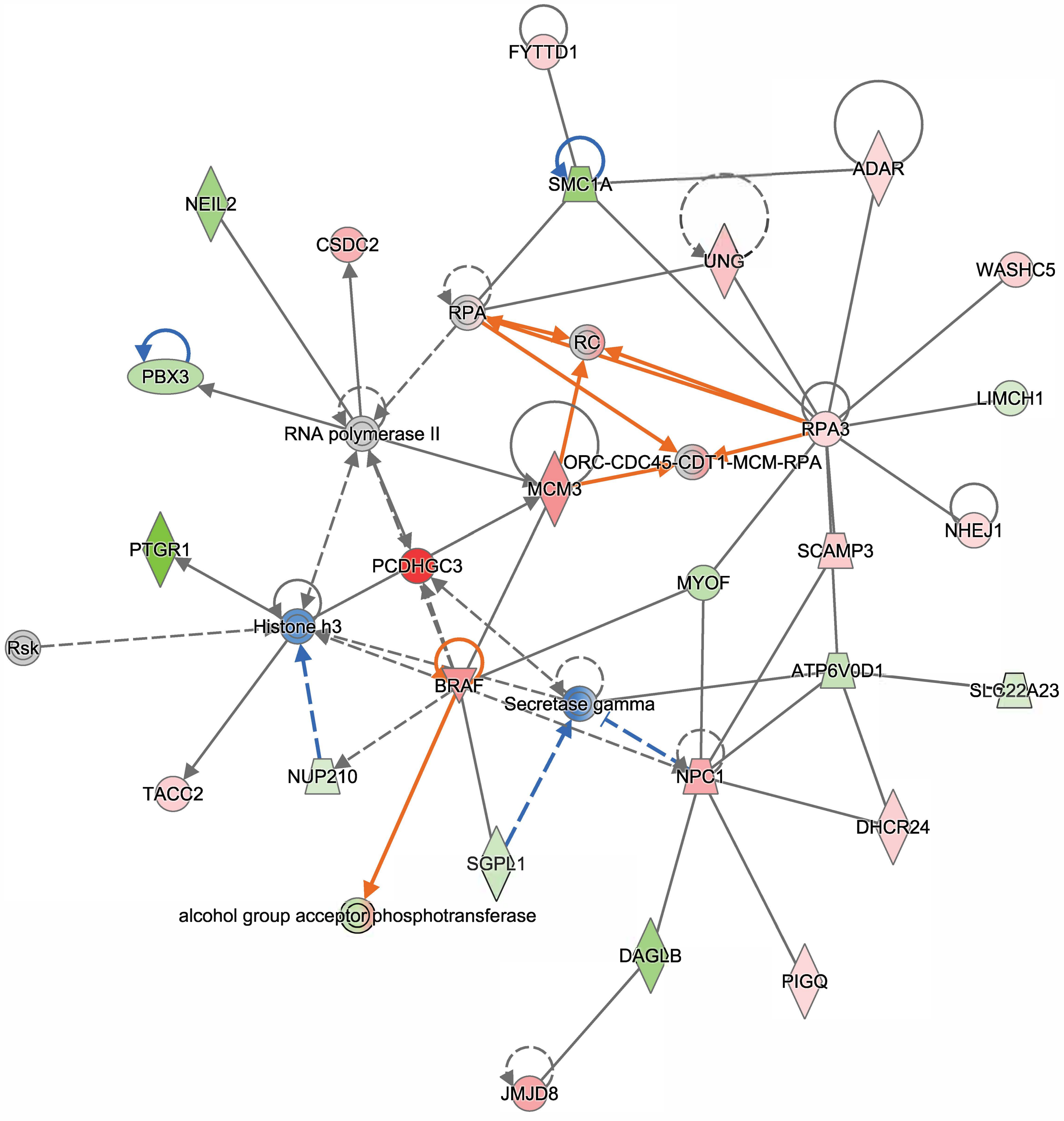


© 2000-2022 QIAGEN. All rights reserved.

**Supplemental Figure 7**
